# Faecal carriage of ESBL-producing *Escherichia coli* in a remote region of Niger

**DOI:** 10.1101/2022.07.07.499258

**Authors:** Hervé Jacquier, Bachir Assao, Françoise Chau, Ousmane Guindo, Bénédicte Condamine, Mélanie Magnan, Antoine Bridier-Nahmias, Nathan Sayingoza-Makombe, Aissatou Moumouni, Anne-Laure Page, Céline Langendorf, Matthew E Coldiron, Erick Denamur, Victoire de Lastours

## Abstract

**Objective:** Whole genome sequencing (WGS) of extended-spectrum β-lactamase-producing *Escherichia coli* (ESBL-*E. coli*) in developing countries is lacking. Here we describe the population structure and molecular characteristics of ESBL-*E. coli* faecal isolates in rural Southern Niger.

**Methods:** Stools of 383 healthy participants were collected among which 92.4% were ESBL-*E. coli* carriers; 90 of these ESBL-*E. coli* containing stools (109 ESBL-*E. coli* isolates) were further analysed by WGS, using short- and long-reads.

**Results:** Most isolates belonged to the commensalism-adapted phylogroup A (83.5%), with high clonal diversity. The *bla*_CTX-M-15_ gene was the major ESBL determinant (98.1%), chromosome-integrated in approximately 50% of cases, in multiple integration sites. When plasmid-borne, *bla*_CTX-M-15_ was found in IncF (57.4%) and IncY plasmids (26.2%). Closely related plasmids were found in different genetic backgrounds. Genomic environment analysis of *bla*_CTX-M-15_ in closely related strains argued for mobilisation between plasmids or from plasmid to chromosome.

**Conclusions:** Massive prevalence of community faecal carriage of CTX-M-15-producing *E. coli* was observed in a rural region of Niger due to the spread of highly diverse A phylogroup commensalism-adapted clones, with frequent chromosomal integration of *bla*_CTX-M-15_. Plasmid spread was also observed. These data suggest a risk of sustainable implementation of ESBL in community faecal carriage.

## Introduction

Extended-Spectrum β-Lactamase-producing (ESBL) Enterobacterales, and particularly *Escherichia coli* (ESBL-*E. coli*) have spread worldwide and become endemic, especially in low-and middle-income countries ^1–5^. The diffusion of ESBL-*E. coli* is mainly the result of the epidemiological success of *bla*_CTX-M_ genes in the 2000s, which are ESBL-encoding genes often carried by mobile genetic elements such as plasmids. *E. coli* are commensals of the gut of vertebrates, including humans ^6^, and constitute the main reservoir of resistance genes in Enterobacterales, leading to ESBL-*E. coli* dissemination in communities ^7^. *E. coli* are also opportunistic pathogens responsible for severe infections such as bloodstream infections, urinary tract infections and in some cases diarrhea ^8^. Infections with ESBL-*E. coli*present a clinical challenge due to limited therapeutic options, which is especially crucial in low-income countries ^9^. Globally, ESBL-*E. coli* infections represent the biggest burden of bacterial antimicrobial resistance (AMR) morbidity and mortality ^10^.

Prevalence of ESBL-*E. coli* faecal carriage in humans is a good indicator of the global burden of antibacterial resistance in a given community ^11^. In Africa, the few available studies showed a prevalence of ESBL-*E. coli* community faecal carriage ranging from 15% to 92% (higher than in most high income countries) depending on the country, the population studied and the year of stool collection ^2–5,12–14^. However, little is known concerning the molecular epidemiology and evolution of resistance mechanisms in low-income countries ^13,15^. Yet, this level of understanding is essential to unravel the evolutionary mechanisms as well as the transmission mechanisms enabling the success of ESBL*-E. coli* in community settings. It is crucial to determine specific strategies to prevent further dissemination of these resistant bacteria. We previously reported that 92.4% of healthy individuals carried ESBL-*E. coli* carried in their stools as part of a trial of ciprofloxacin prophylaxis for meningococcal meningitis performed in a rural region of South Niger ^14^. Faced with such a high prevalence, we sought to describe the population structure of the strains and the molecular basis of this resistance to understand the mechanisms of resistance spread.

## Patients and Methods

### Study design

This work was an ancillary study of a clinical trial on ciprofloxacin prophylaxis against bacterial meningitis carried out between April 22, 2017 and May 18, 2017 during a meningococcal meningitis outbreak in the Madarounfa district in rural southern Niger near the border with Nigeria ^14^.

### Population characteristics

Briefly, a subset of 10 villages in the control arm and 10 villages randomized to receive village-wide prophylaxis in the primary trial were randomly selected to participate in a sub-study examining faecal carriage of ciprofloxacin-resistant and ESBL-producing *Enterobacterales*, at baseline and after interventions ^14^. In each village, 20 households were randomly selected from a census list. One household member in each of the selected households was then selected using a simple lottery. Villages were assigned an identification number and GPS coordinates were collected. Villages were further described by the presence/absence of a market or a health center, as well as the distance to the nearest paved road.

### Strain collection

In the primary trial, stool samples of 383 individuals were collected at day 0 (prior to ciprofloxacin administration in the intervention arm). Samples were collected at home by participants and inoculated into Cary-Blair media (Copan, Brescia, Italy) by study staff. Of them, 99 faecal samples were randomly selected and were sent to the Infection Antimicrobials Modelling and Evolution laboratory (IAME, Université Paris Cité) for quality control in agar deep tubes ^14^. In Paris, faeces were plated on cefotaxime containing plates, and colonies of different morphologies were identified by mass spectrometry (MALDI-TOF-MS Microflex Bruker Daltonics/BioTyper™ version 2.0). When colonies with different morphologies on the same plate were identified as *E. coli*, each isolate was studied independently. Antimicrobial susceptibility testing was performed by the disk diffusion method according to the EUCAST guidelines, and ESBL detection using the synergy test. Among the 99 samples, 90 samples from 90 subjects carrying at least one confirmed ESBL-*E. coli*, constitute the material for the analyses performed in this work.

### Global genomic analysis

For each *E. coli* isolate, DNA was extracted using NucleoMag DNA Bacteria for DNA purification (Macherey Nagel, Hoerdt, France) and the whole genome was sequenced using Illumina Hiseq 2×100 bp after NextEra XT library preparation (Illumina, San Diego, CA, USA).

Sequences were analysed with a previously described in-house bioinformatic pipeline ^16^(https://github.com/iame-researchCenter/Petanc). Briefly, *de novo* assembling was performed using SPAdes 3.10.0 ^17^ and genomes were ordered with Ragout after genetic distance estimation using Mash ^18,19^. Genome typing was performed with *in silico* methods including phylogrouping ^20^, multi-locus sequence typing (MLST) (according to the Warwick University and Pasteur Institute)^21^, serotyping ^22^, and *fim*H typing ^23^. Plasmid replicons were determined using PlasmidFinder ^24^, and the contig location (plasmid or chromosome) using Plascope ^25^. Resistome was analyzed with Resfinder v3.2 ^26^, and the virulome with a database based on Virulence Finder ^27^ and VFDB ^28^. The main virulence factors associated with extra-intestinal or intra-intestinal pathogenic *E. coli* (respectively ExPEC and InPEC) were as in ^29^.

A core genome was determined using Roary v3.12 with default parameters ^30^, and a SNP distance matrix was computed from the core genome (available in Supplementary Table 1). Then, a phylogeny based on a maximum likelihood approach using a core genome alignment was performed using Iqtree v1.6.12 ^31^. Patristic distances obtained were used to design the phylogenetic tree with Itol (v6.3)^32^.

### ESBL-encoded gene support determination

For strains with ESBL-encoded genes embedded in chromosome, the integration sites of *bla*_CTX-M-15_ was mapped on the linkage map of *E. coli* K-12^33^, a A phylogroup strain ST10, assuming synteny wih the studied isolates. Coordinates established by the completed sequence were expressed as 100 minutes for the entire circular map^33^.

For strains with ESBL-encoded genes embedded in plasmids, an additional long-read sequencing was performed. One representative isolate was selected for isolates with close genetic distances according to Mash. The Nanopore sequencing (Oxford, UK) was performed using Ligation Sequencing Kit (SQK-LSK109) with Native Barcoding Expansion 1-12 or 13-24, a FLO--min106 R9.4.1 flowcell, and a MinION device Mk1B. The reads were filtered using Filtlong (v0.2.0) (https://github.com/rrwick/Filtlong) to keep reads longer than 1000 nucleotides. The selected reads were then assembled with the combination of minimap2 (v2.15) and miniasm (v0.3)^34^. The resulting assembly was then polished through 2 rounds of minipolish (v0.1.0), a convenient wrapper for Racon (V1.4.11)^35,36^. llumina and nanopore contigs were subsequently assembled using Unicycler^37^. Plasmids were compared with BRIG^38^ when a scaffold was already available, or circlize R-package^39^ in absence of previously described reference. Plasmids with less than 1% difference in the nucleotide sequences or ≤2 large insertions or deletions (indels) were considered as highly similar. Genome annotation was performed using RAST^40^.

### Ethics

The parent study protocol, which included this sub-study, was reviewed and approved by the National Consultative Ethics Committee of Niger (Ref: 003/2016/CCNE) and the Ethics Review Board of Médecins Sans Frontières (Ref: 1603)^14^. The trial is registered at ClinicalTrials.gov (NCT02724046). Written informed consent was obtained from individual participants.

## Results

All raw data concerning antibiotic resistance and genome typing are available in Supplementary Table 2.

### Population characteristics

In this ancillary study, the median age of the 90 participants was 10.5 years [IQR 4-24.8], and the F:M sex-ratio was 1.5. Only 3 subjects (3.3%) reported antibiotic exposure or hospitalization in the three months prior to sample collection.

### ESBL-*E. coli* isolation and resistance to antibiotics

A total of phenotypically distinct 109 ESBL*-E. coli* were recovered from the 90 samples. Seventy-two participants carried only one ESBL-*E. coli*, 17 carried two distinct ESBL-*E. coli* and one participant carried 3 distinct ESBL-*E. coli* by colony morphology (see above).

Among the 109 ESBL-*E. coli* isolates, 58 (53.2%) were resistant to nalidixic acid, 56 (51.4%) to ciprofloxacin, 23 (21.1%) to gentamicin, 76 (69.7%) to cotrimoxazole, and 85 (78.0%) to tetracycline. All were susceptible to amikacin and imipenem.

### Genomic diversity of ESBL-*E. coli*

The genomic diversity of ESBL-*E. coli* is shown in Table 1 and Figure 1. The ESBL-*E. coli* mainly belonged to phylogroup A (91/109, 83.5%), phylogroup B1 (8/109, 7.3%) and phylogroup C (6/109, 5.5%). One single isolate was found in each of phylogroups D, F, B2 and clade I.

**Table 1.**
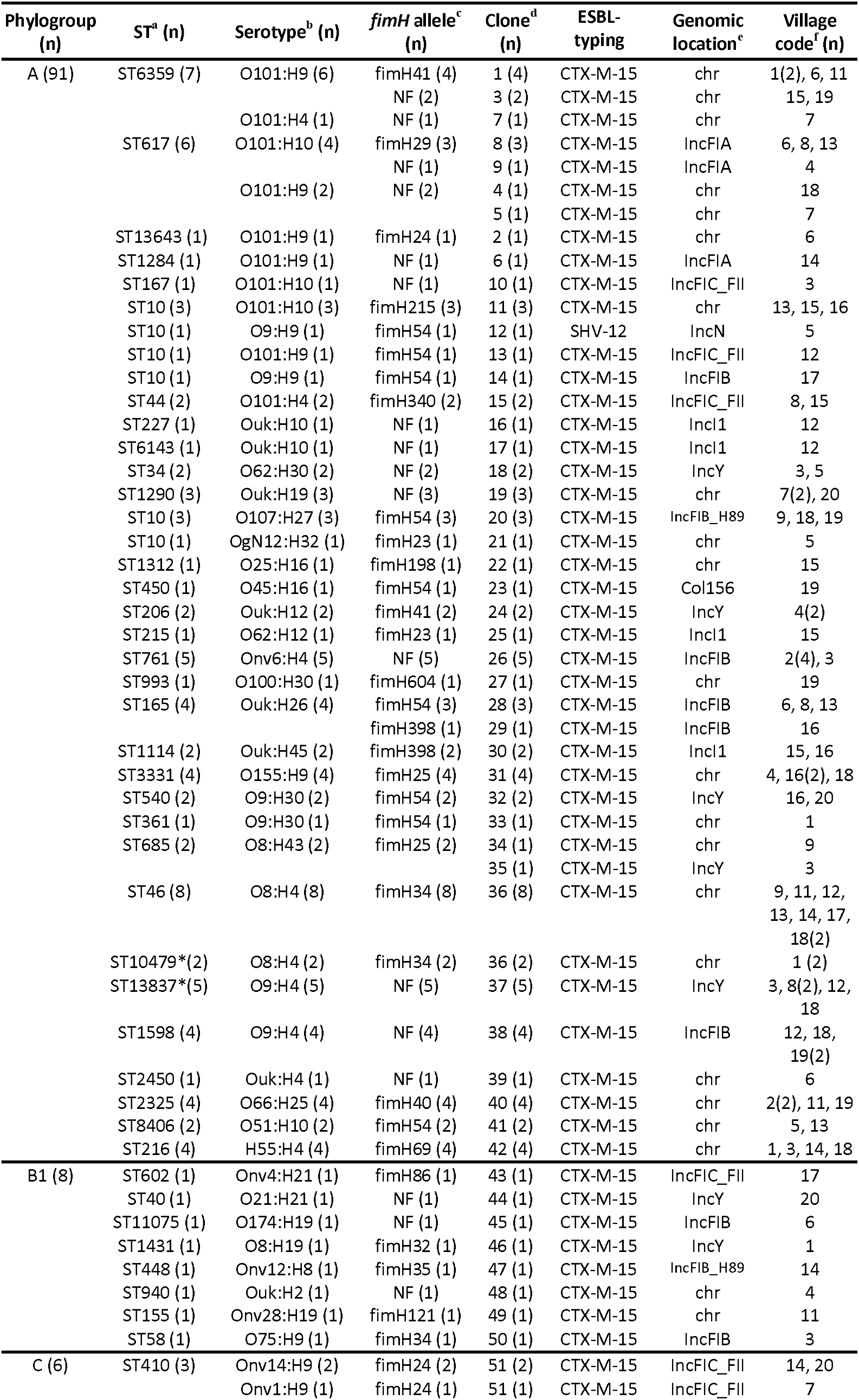
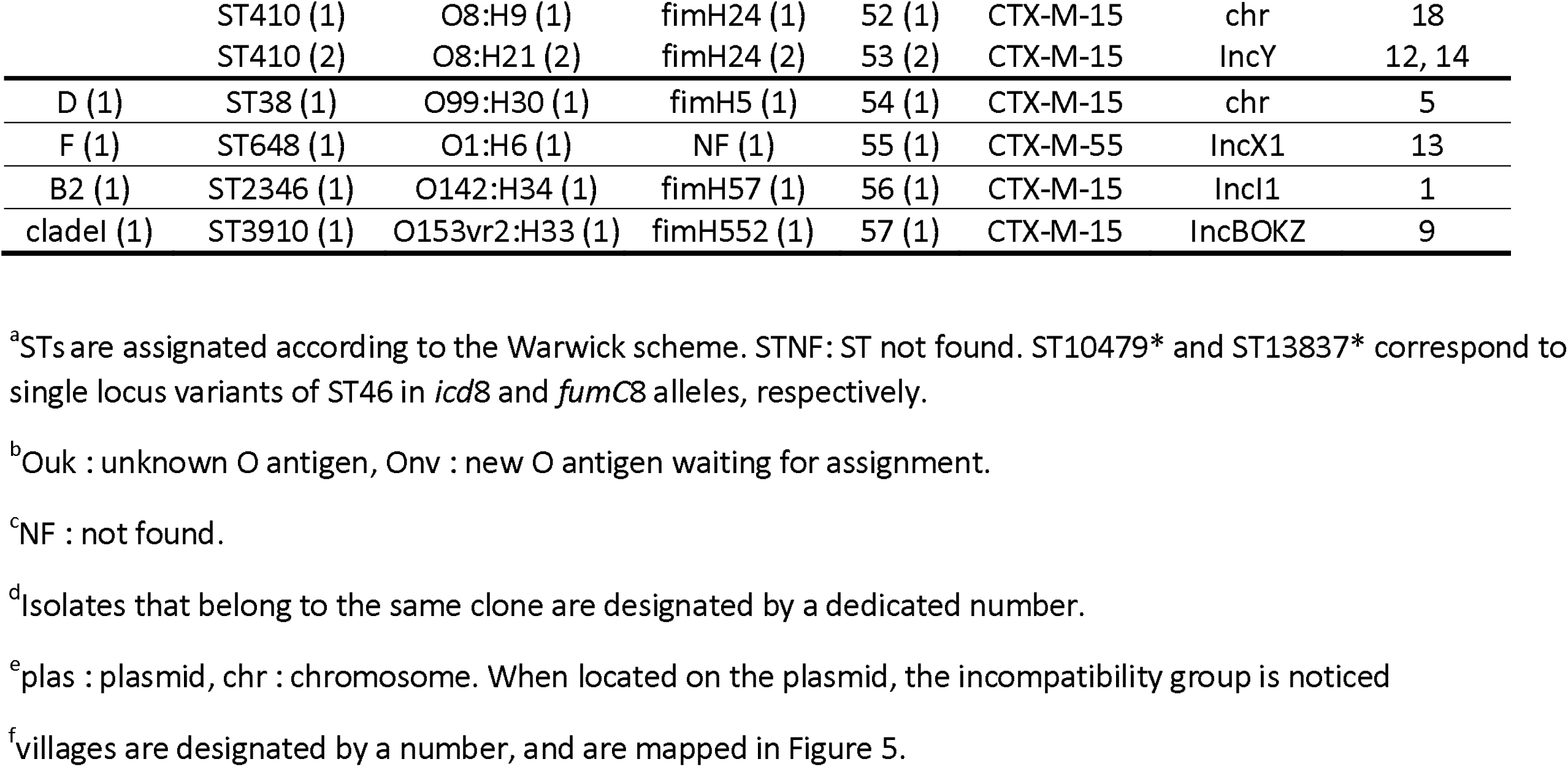
Characteristics of ESBL-*E. coli* according to genome typing and geographical location

**Figure 1.**
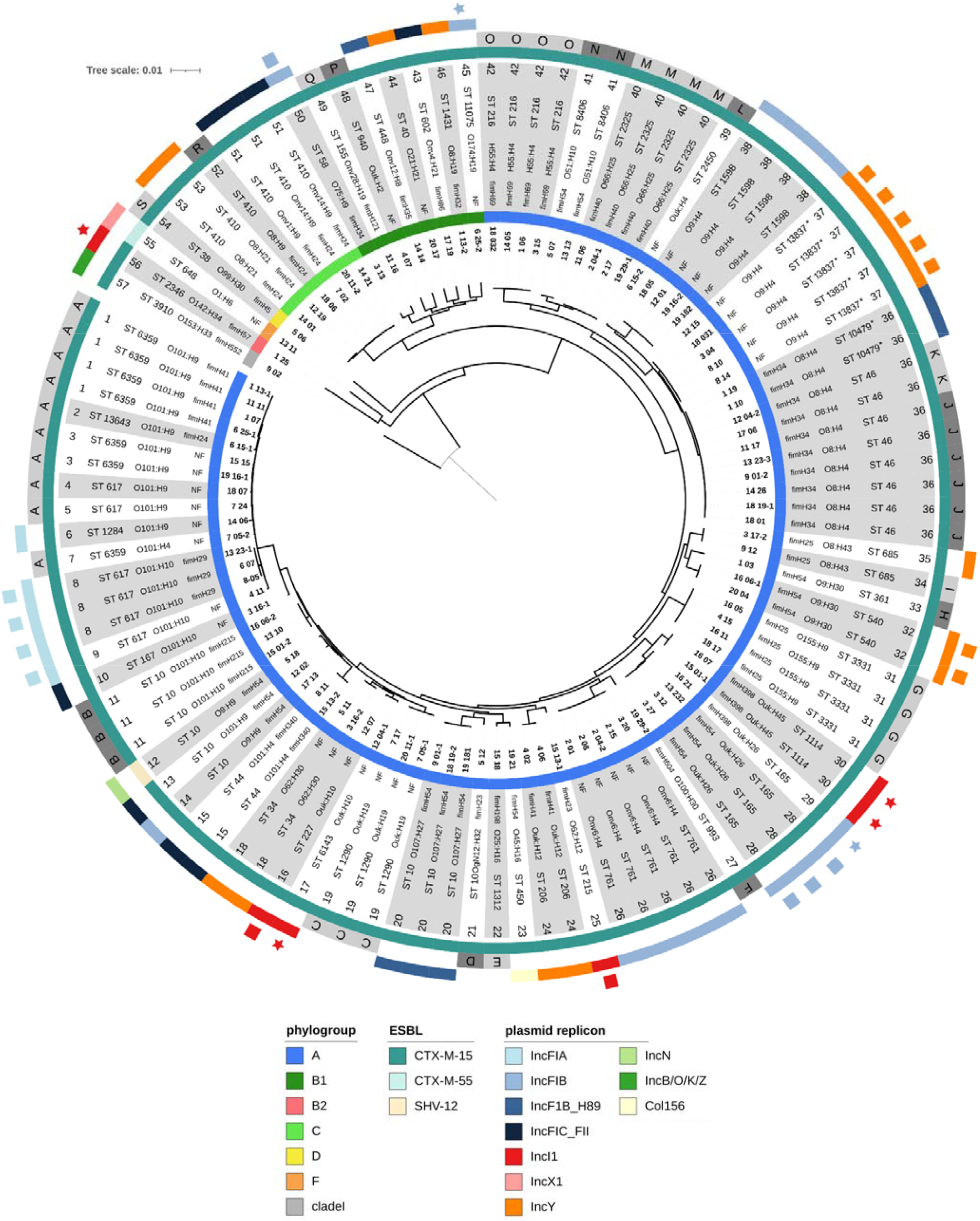
Core genome SNP-based phylogenetic tree of 109 ESBL-*E. coli*. The tree is rooted on *E. coli* clade I (isolate 9_02 (clone 47)). From center to periphery, the different layers correspond to the isolate name, the phylogenetic group, the *fimH* allele, the serotype, the sequence type (according the Warwick University scheme), the clone number, the ESBL-type, the chromosome integration site, and the CTX-M carrying plasmid replicon type. In the layer dedicated to ST type, ST10479 and ST13837 (clones 36 and 37) are single locus variant of ST46 and annotated with a star Squares and stars are used alternatively to separate isolates. Chromosome integration sites (A to S) are described in Supplementary Table 3.

ST typing using Warwick University scheme highlighted 44 different STs. The principal STs identified were ST46 (15/109, 13.8%), ST10 (10/109, 9.2%), ST6359 (7/109, 6.4%), ST617 (6/109, 5.5%), and ST410 (6/109, 5.5%). *In silico* serotyping approaches identified 42 different serotypes, including 11 serotypes with unknown O antigens and new O antigens waiting for assignment for 24/109 (22%) isolates. The main serotypes were O101:H9 (11/109, 10.1%), O101:H4 (10/109, 9.1%), O101:H10 (9/109, 8.3%), and O9:H9 (8/109, 7.3%). A total of 21 different *fimH* alleles were identified. For 34/109 (31.2%) isolates, no *fimH* gene was found. The main *fimH* alleles identified belonged to *fimH54* (15/109, 13.8%), *fimH34* (11/109, 10.1%), *fimH24* (7/109, 6.4%), and *fimH41* (6/109, 5.5%).

### Clone definition and global epidemiology

The definition of clone within the *E. coli* species is not trivial and has been refined at each technological advance from serotype, MLST, *fimH* typing, to more recently whole genome sequencing. The ST entity is heterogeneous in terms of divergence, and can encompass several clones, as exemplified by the emerging ST131 with clades A, B and C having high levels of diversity^8^. Here, the clones were defined according the following strategy. First, isolates were grouped by their haplogroup, defined as a combination of their sequence types (ST) according to the Warwick University and Institute Pasteur schemes, their serotype (O:H), and their *fimH* allele. Second, the distribution of SNPs among isolates was determined, and displayed a SNP cut-off around 120 where few values were observed (Supplementary Figure 1). Third, SNP distances within and between haplogroups were calculated and confronted to the cut-off. By using a cut-off at 120 SNPs, most isolates belonging to a given haplogroup had a SNP distance below this cut-off (Supplementary Figure 1A). A diversity > 120 SNPs was found for few isolates belonging to a given haplogroup: clones 4/5 (683 SNPs), clones 12/14 (2565 SNPs) and clones 34/35 (173 SNPs)(Figure 1). Conversely, the clone 36 was associated with two different haplogroups, due to differences in ST assignation. It should be noted that strain ST10479 is a single locus variant of strain ST46, highlighting a high degree of similarity. Similar results have been found with patristic distances (Supplementary Figure 1B). We thus defined the clone as encompassing isolates with less than 120 SNPs on their core genomes. Based on this definition, 57 distinct clones were identified among the 109 isolates. Of 109 isolates, 74 (67.9%) belonged to clones represented by two or more isolates. Isolates mainly belonged to clones 36 (10/109, 9.2%), 37 (5/109, 4.6%) and 26 (5/109, 4.6%). Of note, *E. coli* strains with different morphologies from the same individual were always genetically distinct (Table1, Figure 1).

In summary, despite belonging almost uniquely to phylogroup A, a huge genomic diversity of clones was observed among the ESBL-*E. coli* isolates.

### ESBL-encoding genes and genomic location

The ESBL phenotype was conferred by the *bla*_CTX-M-15_ gene for 107/109 (98.1%) isolates. The *bla*_CTX-M-15_ gene was located on the chromosome for 48/107 (44.9%) isolates and on a plasmid for 59/107 (55.1%) isolates. The remaining two isolates had the ESBL phenotype conferred by *bla*_SHV-2a_ (1 isolate) and *bla*_CTX-M-55_ genes (1 isolate), and were both located on a plasmid.

The 48 isolates with a chromosomal *bla*_CTXM-15_ gene were phylogenetically diverse, belonging to 23 clones. We identified 19 different chromosome integration sites (Figure 1 and Supplementary Table 3). For most isolates, a given chromosome integration site was associated with a given clone (Figure 1). For a single clone (clone 36), we observed two different integration sites (Supplementary Table 3). Altogether, these data indicate multiple integration events in diverse genetic backgrounds. The distribution of integration sites on the chromosome of *E. coli* K-12 (phylogroup A) has been mapped, assuming global synteny between studied isolates and K-12 (Figure 2). Integration sites have been found in different chromosomal regions, but are not evenly distributed across the genome. Most of the integration sites are within the Ter macrodomain whereas no integration has been identified in the Ori macrodomain ^41^ For isolates with ESBL-encoding gene located on a plasmid, 10 different replicon types were identified. The plasmids with *bla*_SHV-2a_ and *bla*_CTX-M-55_ genes belonged to IncN and IncX1 incompatibility groups, respectively. The plasmids harbouring *bla*_CTX-M-15_ belonged mainly to IncF (IncFIA (5/107), IncFIB (16/107), IncFIB (H89 phagemid plasmid) (6/107), IncFIC(FII) (8/107)) and IncY (16/107) and, to a lesser extent, by IncI1 (6/107) incompatibility groups (Table 1, Figure 3).

**Figure 2.**
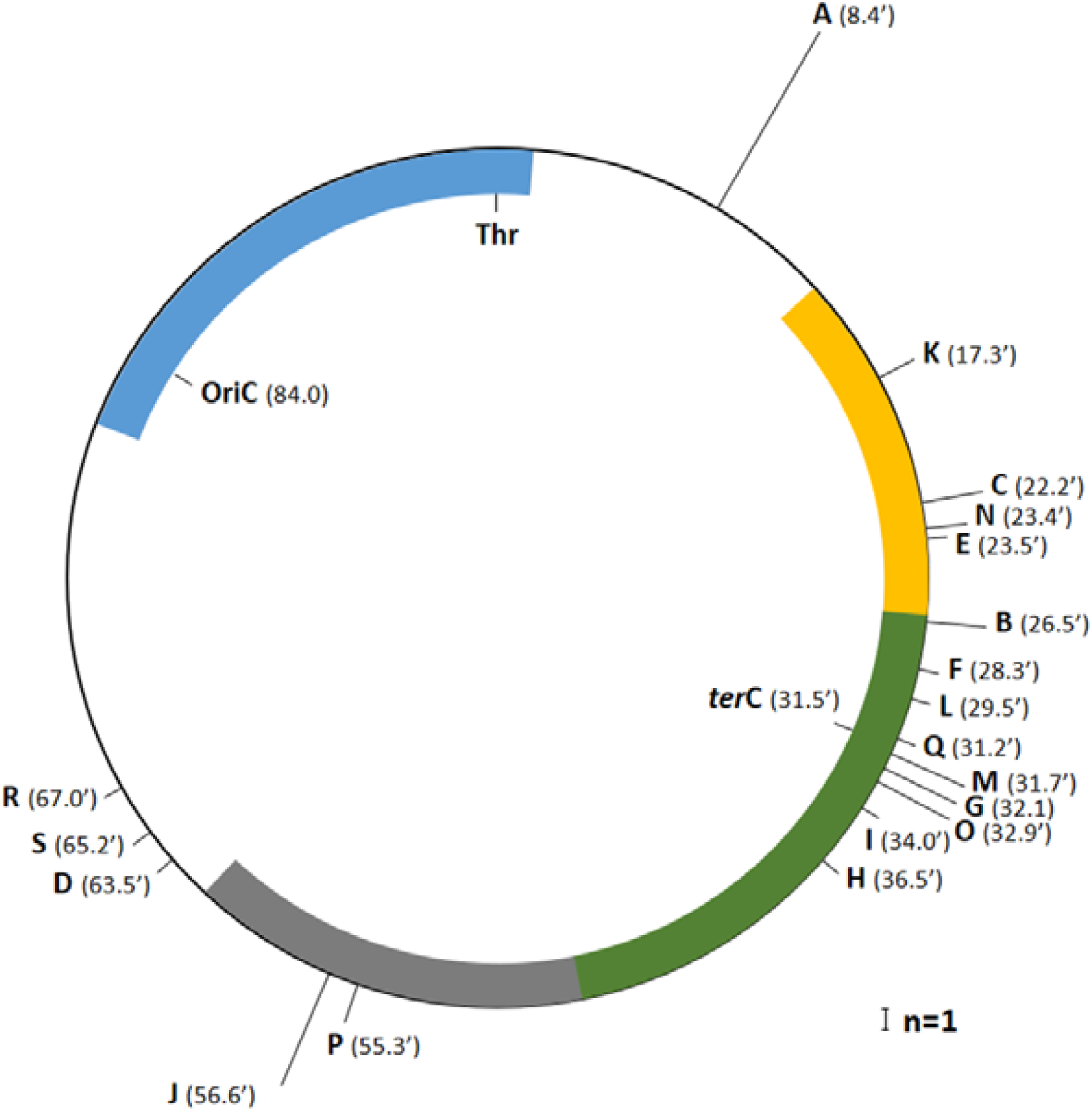
Location of integration sites of *bla*_CTX-M-15_ on the linkage map of *E. coli* K-12. Coordinates established by the completed sequence are expressed as 100 minutes for the entire circular map. The height of each bar is proportional to the number of isolates with a given integration site. Macrodomains are indicated by different colors: Left (13’-26’) in yellow, Ter (26’-47’) in green, Right (47’-62’) in grey and Ori (81’-1’) in blue, as described by Valens *et al*.^41^.

**Figure 3.**
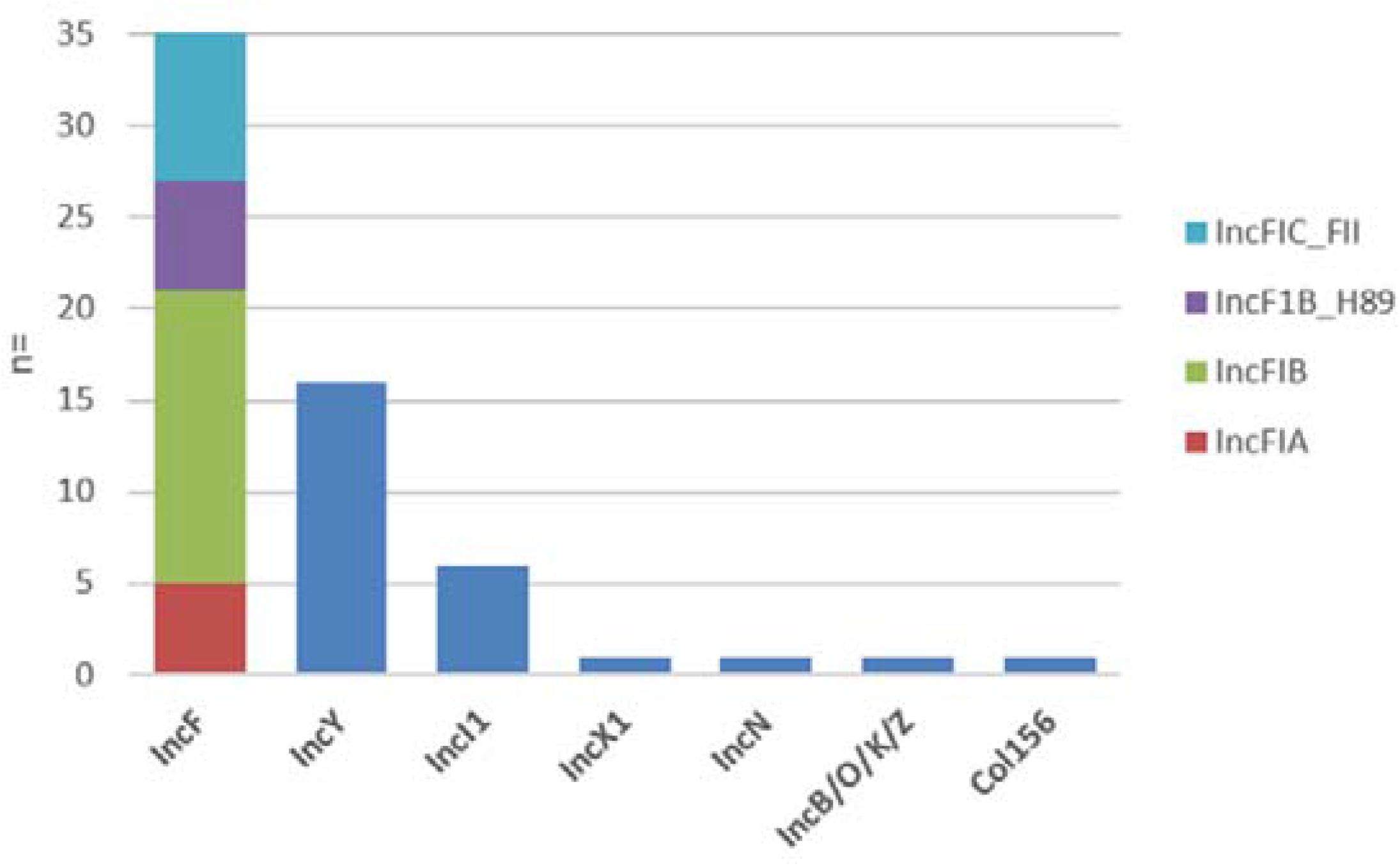
Prevalence of plasmid replicons harboring ESBL-encoding genes.

To better understand the relationships between the clones and the plasmids, we studied the isolates for which the plasmids showed a strong similarity (see above). In six cases, we found highly similar plasmids in different genetic backgrounds, whereas 20 plasmids were strictly associated with a single genetic background (for plasmid alignments, see Supplementary Figure 2). For incFIA plasmids, we identified similar plasmids, with insertion of 26.7 kbp in the plasmid hosted by clone 8 compared to that of clone 9. For IncFIB plasmids, we identified similar plasmids in clones 28 and 50, with insertion of 26.7 kbp in the plasmid hosted by clone 28, and an insertion of 8.8 kbp in the plasmid hosted by clone 50. Moreover, the plasmid hosted by clone 29 presented a 27.9 kbp insertion compared to that of clone 45. For IncI1 plasmids, plasmids hosted by clones 16 and 25 presented more than 99% identity, and the ones hosted by the clones 17, 30, and 56 presented two indels of 2.14 and 1.34 kbp. For IncY plasmids, we identified similar plasmids, with insertion of 17 kbp in the plasmid hosted by clone 32 compared to that of clone 37. Altogether, these data suggest plasmid transmission between clones.

### Focus on evolutionary scenarios of *bla*_CTX-M-15_ gene spread in closely related clones

If our analyses suggest both clonal and plasmid spreads, we wanted to go further in the unravelling of putative evolutionary scenarios leading to the diffusion of *bla*_CTX-M-15_ gene in closely related isolates. Here we focus on two clusters: one with clones 1 to 7, and a second with clones 36 and 37.

### Focus on clones 1 to 7

For isolates with *bla*_CTX-M-15_ gene embedded in the integration site “A” (clones 1 to 5 and 7), we assume that *bla*_CTX-M-15_ gene acquisition would have followed the most parsimonious scenario: isolates with chromosomal *bla*_CTX-M-15_ gene have diversified after a single event of integration in a common ancestor. This assumption is based on the fact that the insertion site was the same for all isolates with chromosomal *bla*_CTX-M-15_ and that IS*EcpI* upstream *bla*_CTX-M-15_ is largely deleted (data not shown). In addition, in the closely-related clone 6, *bla*_CTX-M-15_ gene was acquired through an IncFIA plasmid.

### *Focus on clones 36* and *37*

Conversely, in clone 36/ST46, *bla*_CTX-M-15_ is hosted in two different chromosome integration sites, J (six isolates) and K (two isolates), suggesting two independent integration events in the same clone. In addition, in the closely-related clones 36/ST10479 and clone 37, *bla*_CTX-M-15_ gene was acquired by the IncFIB and IncY plasmids, respectively.

Analysis of synteny in the regions flanking the *bla*_CTX-M-15_ gene (Figure 4) displays highly conserved genetic environment, suggesting transfer between plasmid and chromosome and/or between plasmids.

**Figure 4.**
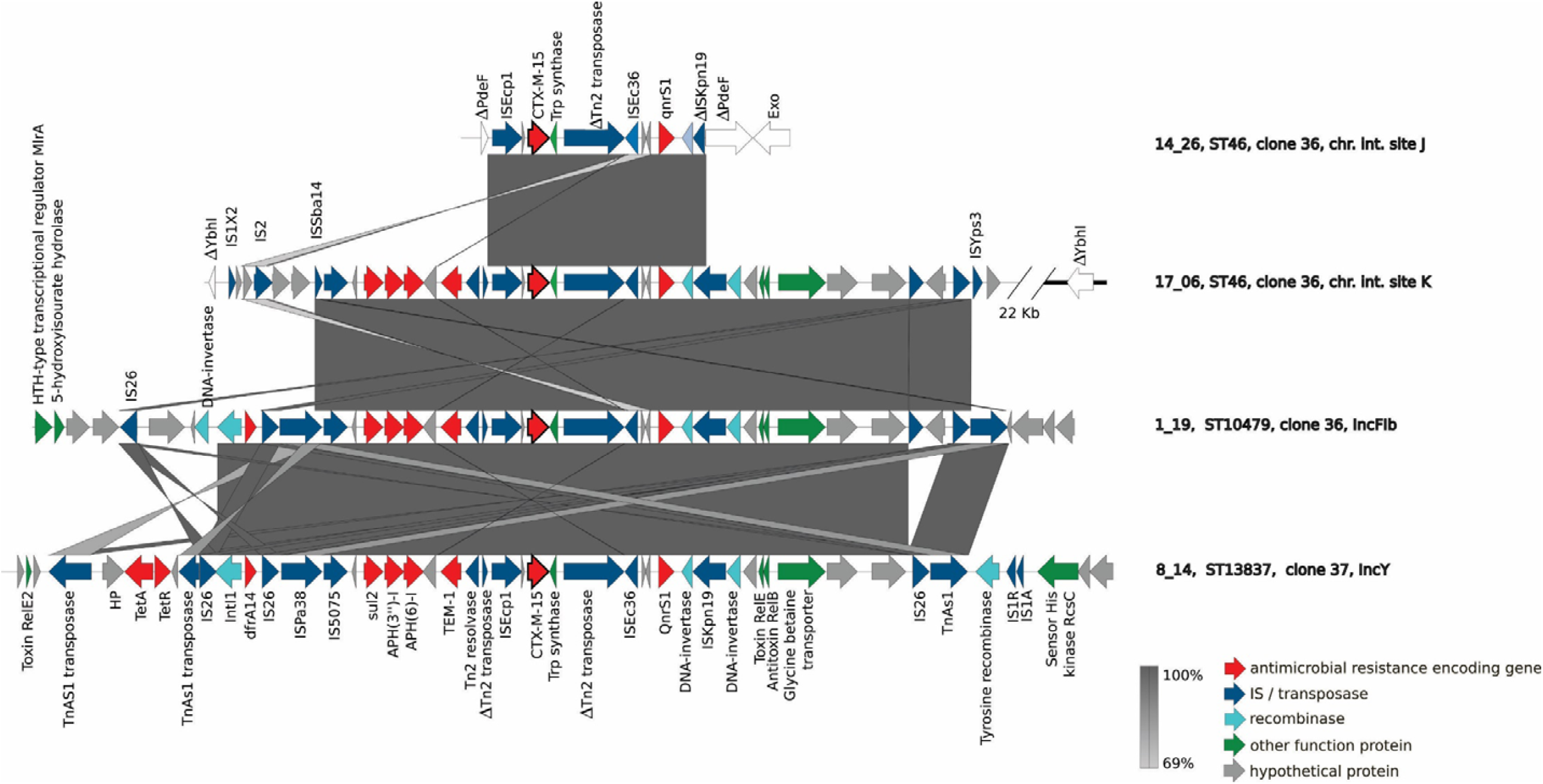
Genetic environments of CTX-M-15 encoding genes for four isolates representative of different genetic patterns of ST46 and ST46 single locus variants For the four isolates, the number of the isolate, the single variant locus of ST46, the clone number, and the chromosome integration site or plasmid type are indicated on the right of the figure.

### Virulence associated genes

We focused on the main virulence factors associated with extra-intestinal or intra-intestinal pathogenic *E. coli* (respectively ExPEC and InPEC)(19). The average number of ExPEC virulence genes was low (1.24), the most prevalent being *fuyA* and *irp2* (25/109, 22.9%) (Supplementary Table 2). These two genes encode the yersiniabactin siderophore and its transporter, and are known to be part of the high pathogenicity island (HPI)^42^. For phylogroup A isolates, these two genes were unevenly distributed among the strains, and were mostly found in a cluster of clones (clones 1 to 6, and clone 8). In the phylogroup C, most isolates (4/6) harboured the adhesin encoding *pap* genes in addition to *fuyA* and *irp2* genes. One enterotoxinogenic *E. coli* (ETEC) (*eltB* positive) from clade I and two enteropathogenic *E. coli* (EPEC) from B1 and B2 phylogroups were identified, the latter belonging to the ST2346 and being a typical EPEC (*eae* and *bfp* positive)^43^.

### Geographical origin of ESBL-*E. coli* carriers

The geographical distribution of the villages of the carriers of the 109 ESBL-*E. coli* is represented in Figure 5. Most isolates belonging to the same clone were not limited to a given geographical area, but were widely spread across the different villages (such as clones 1, 8, 15, 20, 36, 37, 40, 41, and 42). Nevertheless, few clones, had a local spread limited to 1 or 2 close villages (such as clones 3, 18, 19, 24, and 26). Last, some clones widely spread in the eastern area, in villages connected by a paved road and which depend on the same healthcare center (clones 11, 30, 31, 32, 38, 51, and 53).

**Figure 2.**
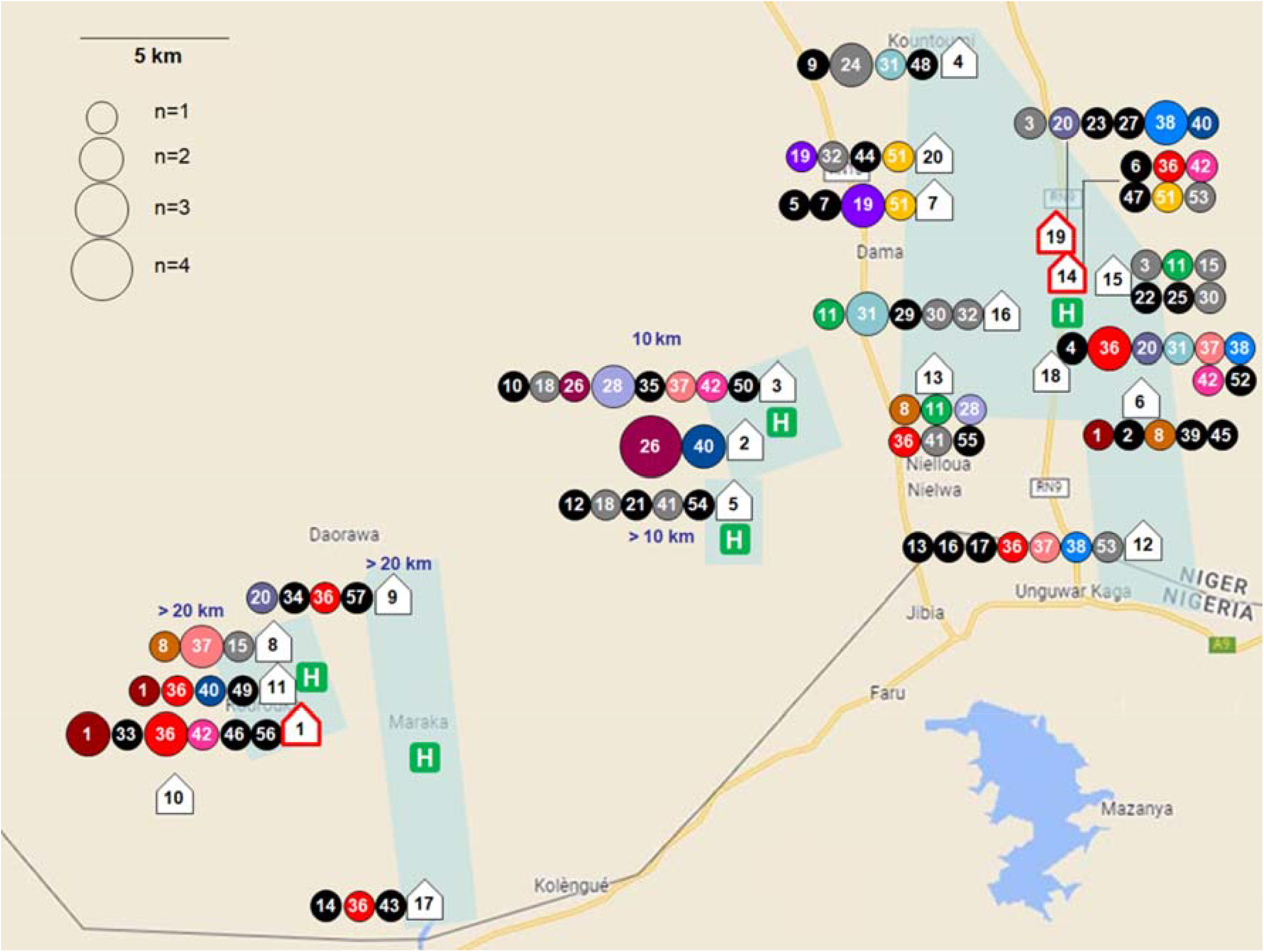
Geographical location of different clones of ESBL-*E. coli* House icons represent the locations of the villages. No isolates were recovered from village 10. Villages with a market are bordered in red. Healthcare centres are indicated by a green box with an H. Villages belonging to the same healthcare district appear in the light blue shapes. Distance to the nearest paved road are indicated in blue. For each village, circles represent clones whose area is proportional to the number of isolates. Circles coloured with different colours correspond to clones represented by more than two isolates, those in grey and black to clones represented by two or one isolate, respectively.

The isolates 3_13 (belonging to clone 50) and the isolates 3_12 and 3_27 (belonging to clone 28) harbour plasmids with strong similarity, and were identified in the same village (village 3), suggesting a putative plasmid transfer between isolates with different genetic backgrounds (Figure 1 and Figure 4).

In sum, we did not find any correlation between geographical localization of the subjects and the epidemiological genomic data of ESBL-*E. coli*, which suggests a large spread of the most frequent clones.

## Discussion

After finding an unexpectedly high prevalence of ESBL-*E. coli* faecal carriage in healthy subjects (>90%) in rural South Niger ^14^, we analysed the population structure, molecular support of resistance and mode of spread of the resistance in these commensal strains, using state-of-the-art sequencing techniques with both Illumina WGS and an additional long-read sequencing by Nanopore to specifically study strains with ESBL-encoded genes embedded in plasmids. Previous studies of ESBL carriage in African settings has varied by the region, the community and the age of the subjects ^2–5,12,13^. Reported prevalence in Africa has been high, ranging from 46 % in district hospital patients in South Africa in 2018 ^3^, 51% in hospitalized patients and 38% in healthy healthcare workers in Chad in 2019 ^13^, and 59% in healthy children in the community in the Central African Republic ^12^. When resistance mechanisms and phylogenetic groups were described, *bla*_CTX-M-15_ was the most commonly encountered gene encoding for resistance to β-lactams (ranging from 75% to 90% of cases), and the clonal group ST131 was predominant among phylogroup B2 *E. coli* ^12,13,15,44^.

The main features of our work are the following. First, most ESBL-*E. coli* belonged to phylogroup A (83.4%), but with a high diversity in terms of STs, serotypes, *fimH* alleles and finally clones, even though some clones were overrepresented. *E. coli* strains belonging to phylogroup A are known to be adapted to human gut commensalism lifestyle ^6^ and have been described as predominant in low-and middle-income countries ^45^. As expected, these strains carry few virulence genes, with the exception of a cluster of clones (clones 1 to 6, and clone 8), which have accumulated many virulence genes, especially iron capture systems such as yersiniabactin (*fyuA, irp2*), borne on the HPI (Supplementary Table 2). ST410 (phylogroup C), an emerging high-risk clone ^46^, is also well represented, with strains harbouring the *pap* adhesin encoding genes in addition to the HPI. Both gene clusters have been associated with ESBL-*E. coli* strains with a longer duration of faecal carriage in travellers returning from developing countries ^47^. On the other hand, B2 phylogroup was represented by a single isolate, a typical EPEC belonging to the EPEC9 cluster ^43,48,49^. This is in contrast to what has been observed in Western countries, where the proportion of B2 strains among commensal *E. coli* has dramatically increased in the last four decades, probably under the pressure of several factors such as changes in dietary habits and improved hygiene ^6,50^. Of note, no ST131-*E. coli* isolate (also a B2) was found here, despite the clone’s global epidemiological success ^12,13,51^. The pronounced diversity among A strains and the low prevalence of B2 strains supports the hypothesis that the strains described here are almost individual-specific, and aligns with the expected composition of commensal *E. coli* strains from humans living in developing countries. Along the same line, a recent study performed in healthy pregnant women in Madagascar showed a high prevalence of ESBL-*E. coli* (34%), composed mostly of isolates belonging to various phylogroup A clones (72%)^52^.

Second, we found that the ESBL phenotype was mainly associated with the *bla*_CTX-M-15_ gene (98%), which was embedded in the chromosome in approximately 50% of cases, at different integration sites. The spread of the *bla*_CTX-M-15_ gene has been well-described in the last decades, even though other *bla*_CTX-M_ alleles such as *bla*_CTX-M-14_ or *bla*_CTX-M-27_ have had some epidemiological success ^2^. As with most AMR genes, the spread of *bla*_CTX-M_ genes has been associated with plasmid dissemination, particularly the pC15-1a plasmid ^53^. However, chromosomal locations have increasingly been described. Chromosome-located *bla*_CTX-M_ genes were found in 4.8% of ESBL-*E. coli* isolates in a veterinary collection from Germany, the Netherlands, and the UK ^54^. This rate increases to 8.3% and 14.6%, respectively, in clinical isolates recovered from hospitals in Zambia ^55^ and China ^56^. In the study from Madagascar cited above ^52^, the *bla* gene was located on the chromosome in 31% of strains. The highest incidence of chromosomal *bla*_CTX-M_ gene described so far was observed in Japan, where *bla*_CTX-M-14_ was found on the chromosome in 45.9% of cases, and in 16.2% both on chromosome and plasmid in community commensal strains ^57^. The location of *bla*_CTX-M_ genes on both chromosome and plasmid probably reflects the ability of their genetic environment to mobilize these resistance genes. In their model, Hamamoto *et al*. display high rates of *bla*_CTX-M-14_ mobilization, and integration in various sites ^58^, as was observed here with our isolates. Whether a chromosomal or plasmid location confers a selective advantage for the resistance genes is a matter of debate. In a mathematical model, Lethtinen *et al*. suggested that plasmid location could be preferred under selective pressure ^59^, suggesting that a chromosomal location is more adapted to a lower levels of selective pressure, as would be the case in community carriage. This hypothesis is also supported by the fitness cost associated with plasmid-carried resistance genes ^60^ and by the conservation of vertical transfer of chromosome-located genes. These assumptions remain challenged by other authors, as compensatory mutations alleviating plasmid fitness costs and high plasmid-chromosome beneficial interactions have also been described ^60^. From an epidemiological point-of-view, a recent work suggested that chromosome integration of *bla*_CTX-M-15_ in ST131 isolates could have contributed to the clonal expansion of a rare C2 lineage ^61^.

Third, for isolates with plasmid-located *bla*_CTX-M-15_, we showed a predominance of IncF plasmids, known to be well adapted to *Enterobacterales*, followed by IncY plasmids, also reported in Madagascar. Interestingly, a high similarity was found for some plasmids in different genetic backgrounds, suggesting both isolate and plasmid diffusion. This observation here in a human community is in line with a previous study of temporal intestinal carriage of ESBL-*E. coli* in calf farms ^62^. Additionally, we found a similarity between IncFIb plasmid backbones and pCFSAN061762, a plasmid described in Egypt in an isolate of *E. coli* ST38 (data not shown).

Fourth, careful genetic analysis of the *bla*_CTX-M-15_ gene in the ST46 and ST46 single locus variant clones (Figure 4) argues for mobilisation of the ESBL gene with variable numbers of flanking genes between plasmids and from plasmids to chromosome. This process has been recently suggested in A phylogroup strains isolated during a “One Health” study on Reunion island ^63^.

Finally, we did not find any correlation between geographical localization of the subjects (the villages they live in or their usual social interactions, such as marketplaces) and the epidemiological genomic data of ESBL-*E. coli*, which suggests a large spread of the most frequent clones. These results may unveil an underestimated mix of populations in this area, or contamination by a common source of food or water that could not be captured by mapping health centers, distance to an asphalt road or marketplaces, as represented in Figure 4.

Our work has some limitations: (i) only a subset of stool samples from the overall study was analysed, potentially introducing biases, yet the random selection should have reduced this risk; (ii) only the predominant cefotaxime-resistant *E. coli* isolates present in the faeces were studied, which may not represent the diversity of *E. coli* isolates in the gut, but does represent the quantitatively predominant *E. coli* isolates and therefore the ones that constitute the biggest threat in terms of infectious risks and dissemination ^64^; (iii) the search of an ESBL*-E. coli* reservoir in other environments, such as the food chain (poultry/livestock), water or sediments has not been planned, impeding the understanding of the routes of dissemination in a “One Health” approach ^11^.

Despite these limitations, this epidemiologic study in a remote region of Niger is unique and shows the unprecedented dissemination of ESBL-*E. coli* in the gut of a geographically localized human population. It suggests the widespread dissemination of multiple resistant clones, possibly fostered by underestimated inadequate antibiotic treatments and multiple population exchanges that have probably lasted for years already. From a molecular point of view, this work highlights the diversity of pathways followed by *bla*_CTX-M-15_. The high frequency of chromosomal *bla*_CTX-M-15_ in isolates of *E. coli* belonging to the commensal phylogroup A is of particular concern because it could be associated with their sustainable implementation in the community. Understanding the evolutionary forces that select for such a high prevalence of ESBL-*E. coli* should be major objective.

Altogether, antibiotic resistance in commensal *E. coli* in this remote part of Africa appears to have all the characteristics of long-term adaptation and all the tools for further dissemination. This is especially worrying as commensal *E. coli* are responsible for clinical infections and antimicrobial resistant *E. coli* infections represent the biggest threat in terms of AMR globally ^10^. Noteworthy, community-acquired bacteremia in children hospitalized for severe acute malnutrition was recently followed in an intensive nutritional rehabilitation center in the same rural district. Approximately two-thirds of the *E. coli* and *K. pneumoniae* strains (16/26, 61.5%) were ESBL producers, reinforcing the link between the widespread carriage of ESBL-producing *E. coli* strains and the risk of severe infection from these same microorganisms ^65^. Infections with these strains will be increasingly difficult to treat, with the need for broader spectrum antibiotics, which in turn may select more resistant strains such as carbapenemase producers. This work may just be exposing the tip of the iceberg of an AMR epidemiological catastrophe well on its way.

## Acknowledgments

We offer sincere thanks to all the participants of the study, as well as the staff from Maradi laboratory for their excellent fieldwork, both in terms of sample collection and phenotypic microbiological analyses. We thank “Médecins Sans Frontières” teams for their trust and involvement in the parent study as well as this ancillary study. We thank Epicentre and Rebecca Grais, for trusting our laboratory for the quality control and genetic analyses from the beginning of this work and for the opportunity to gather an impressive collection of faecal samples from such a remote region. We are grateful to Olivier Clermont for his expertise in the analysis of the recently annotated STs.

## Financial support

This work was partially supported by a grant from the “Fondation pour la Recherche Médicale” (Equipe FRM 2016, grant number DEQ20161136698). The parent study was funded by “Médecins Sans Frontières”.

## Declaration of interest statement

None

## Notes

### Competing Interest Statement

The authors have declared no competing interest.

### Summary of Updates

- The definition of clone has been clarified - The following data have been added to the modified manuscript: *mapping of integration sites on the E. coli K12 chromosome *figure with plasmid type distribution *figure with genetic environments of CTX-M-15 encoding genes for isolates representative of different genetic patterns of ST46 and ST46 single locus variants

## References

1 Karanika Styliani, Karantanos Theodoros, Arvanitis Marios, Grigoras Christos, Mylonakis Eleftherios. Fecal Colonization With Extended-spectrum Beta-lactamase–Producing Enterobacteriaceae and Risk Factors Among Healthy Individuals: A Systematic Review and Metaanalysis. Clin Infect Dis 2016;63(3):310–8. Doi: 10.1093/cid/ciw283.

2 Bevan Edward R., Jones Annie M., Hawkey Peter M. Global epidemiology of CTX-M β-lactamases: temporal and geographical shifts in genotype. J Antimicrob Chemother 2017;72(8):2145–55. Doi: 10.1093/jac/dkx146.

3 Founou Raspail Carrel, Founou Luria Leslie, Essack Sabiha Yusuf. Extended spectrum beta-lactamase mediated resistance in carriage and clinical gram-negative ESKAPE bacteria: a comparative study between a district and tertiary hospital in South Africa. Antimicrob Resist Infect Control 2018;7:134. Doi: 10.1186/s13756-018-0423-0.

4 Letara Nuru, Ngocho James Samwel, Karami Nahid, Msuya Sia E., Nyombi Balthazar, Kassam Nancy A., et al. Prevalence and patient related factors associated with Extended-Spectrum Beta-Lactamase producing Escherichia coli and Klebsiella pneumoniae carriage and infection among pediatric patients in Tanzania. Sci Rep 2021;11(1):22759. Doi: 10.1038/s41598-021-02186-2.

5 Diriba Kuma, Awulachew Ephrem, Tekele Lami, Ashuro Zemachu. Fecal Carriage Rate of Extended-Spectrum Beta-Lactamase-Producing Escherichia coli and Klebsiella pneumoniae Among Apparently Health Food Handlers in Dilla University Student Cafeteria. Infect Drug Resist 2020;13:3791–800. Doi: 10.2147/IDR.S269425.

6 Tenaillon Olivier, Skurnik David, Picard Bertrand, Denamur Erick. The population genetics of commensal Escherichia coli. Nat Rev Microbiol 2010;8(3):207–17. Doi: 10.1038/nrmicro2298.

7 Woerther Paul-Louis, Burdet Charles, Chachaty Elisabeth, Andremont Antoine. Trends in human fecal carriage of extended-spectrum β-lactamases in the community: toward the globalization of CTX-M. Clin Microbiol Rev 2013;26(4):744–58. Doi: 10.1128/CMR.00023-13.

8 Denamur Erick, Clermont Olivier, Bonacorsi Stéphane, Gordon David. The population genetics of pathogenic Escherichia coli. Nat Rev Microbiol 2021;19(1):37–54. Doi: 10.1038/s41579-020-0416-x.

9 Ouedraogo A. S., Jean Pierre H., Bañuls A. L., Ouédraogo R., Godreuil S. Emergence and spread of antibiotic resistance in West Africa: contributing factors and threat assessment. Med Sante Trop 2017;27(2):147–54. Doi: 10.1684/mst.2017.0678.

10 Murray Christopher JL, Ikuta Kevin Shunji, Sharara Fablina, Swetschinski Lucien, Aguilar Gisela Robles, Gray Authia, et al. Global burden of bacterial antimicrobial resistance in 2019: a systematic analysis. The Lancet 2022;399(10325):629–55. Doi: 10.1016/S0140-6736(21)02724-0.

11 Puspandari Nelly, Sunarno Sunarno, Febrianti Tati, Febriyana Dwi, Saraswati Ratih Dian, Rooslamiati Indri, et al. Extended spectrum beta-lactamase-producing Escherichia coli surveillance in the human, food chain, and environment sectors: Tricycle project (pilot) in Indonesia. One Health 2021;13:100331. Doi: 10.1016/j.onehlt.2021.100331.

12 Farra A., Frank T., Tondeur L., Bata P., Gody J. C., Onambele M., et al. High rate of faecal carriage of extended-spectrum β-lactamase-producing Enterobacteriaceae in healthy children in Bangui, Central African Republic. Clin Microbiol Infect 2016;22(10):891.e1-891.e4. Doi: 10.1016/j.cmi.2016.07.001.

13 Ouchar Mahamat Oumar, Tidjani Abdelsalam, Lounnas Manon, Hide Mallorie, Benavides Julio, Somasse Calèbe, et al. Fecal carriage of extended-spectrum β-lactamase-producing Enterobacteriaceae in hospital and community settings in Chad. Antimicrob Resist Infect Control 2019;8:169. Doi: 10.1186/s13756-019-0626-z.

14 Coldiron Matthew E., Assao Bachir, Page Anne-Laure, Hitchings Matt D. T., Alcoba Gabriel, Ciglenecki Iza, et al. Single-dose oral ciprofloxacin prophylaxis as a response to a meningococcal meningitis epidemic in the African meningitis belt: A 3-arm, open-label, cluster-randomized trial. PLoS Med 2018;15(6):e1002593. Doi: 10.1371/journal.pmed.1002593.

15 Maataoui Naouale, Langendorf Céline, Berthe Fatou, Bayjanov Jumamurat R., van Schaik Willem, Isanaka Sheila, et al. Increased risk of acquisition and transmission of ESBL-producing Enterobacteriaceae in malnourished children exposed to amoxicillin. J Antimicrob Chemother 2020;75(3):709–17. Doi: 10.1093/jac/dkz487.

16 Bourrel Anne Sophie, Poirel Laurent, Royer Guilhem, Darty Mélanie, Vuillemin Xavier, Kieffer Nicolas, et al. Colistin resistance in Parisian inpatient faecal Escherichia coli as the result of two distinct evolutionary pathways. J Antimicrob Chemother 2019;74(6):1521–30. Doi: 10.1093/jac/dkz090.

17 Bankevich Anton, Nurk Sergey, Antipov Dmitry, Gurevich Alexey A., Dvorkin Mikhail, Kulikov Alexander S., et al. SPAdes: A New Genome Assembly Algorithm and Its Applications to Single-Cell Sequencing. J Comput Biol 2012;19(5):455–77. Doi: 10.1089/cmb.2012.0021.

18 Ondov Brian D., Treangen Todd J., Melsted Páll Mallonee Adam B., Bergman Nicholas H., Koren Sergey, et al. Mash: fast genome and metagenome distance estimation using MinHash. Genome Biol 2016;17(1):132. Doi: 10.1186/s13059-016-0997-x.

19 Kolmogorov Mikhail, Raney Brian, Paten Benedict, Pham Son. Ragout-a reference-assisted assembly tool for bacterial genomes. Bioinforma Oxf Engl 2014;30(12):i302–309. Doi: 10.1093/bioinformatics/btu280.

20 Beghain Johann, Bridier-Nahmias Antoine, Le Nagard Hervé, Denamur Erick, Clermont Olivier. ClermonTyping: an easy-to-use and accurate in silico method for Escherichia genus strain phylotyping. Microb Genomics 2018;4(7). Doi: 10.1099/mgen.0.000192.

21 Jolley Keith A., Bray James E., Maiden Martin C. J. Open-access bacterial population genomics: BIGSdb software, the PubMLST.org website and their applications. Wellcome Open Res 2018;3:124. Doi: 10.12688/wellcomeopenres.14826.1.

22 Ingle Danielle J., Valcanis Mary, Kuzevski Alex, Tauschek Marija, Inouye Michael, Stinear Tim, et al. In silico serotyping of E. coli from short read data identifies limited novel O-loci but extensive diversity of O:H serotype combinations within and between pathogenic lineages. Microb Genomics 2016;2(7):e000064. Doi: 10.1099/mgen.0.000064.

23 Roer Louise, Tchesnokova Veronika, Allesøe Rosa, Muradova Mariya, Chattopadhyay Sujay, Ahrenfeldt Johanne, et al. Development of a Web Tool for Escherichia coli Subtyping Based on fimH Alleles. J Clin Microbiol 2017;55(8):2538–43. Doi: 10.1128/JCM.00737-17.

24 Carattoli Alessandra, Zankari Ea, García-Fernández Aurora, Voldby Larsen Mette, Lund Ole, Villa Laura, et al. In silico detection and typing of plasmids using PlasmidFinder and plasmid multilocus sequence typing. Antimicrob Agents Chemother 2014;58(7):3895–903. Doi: 10.1128/AAC.02412-14.

25 Royer G., Decousser J. W., Branger C., Dubois M., Médigue C., Denamur E., et al. PlaScope: a targeted approach to assess the plasmidome from genome assemblies at the species level. Microb Genomics 2018;4(9). Doi: 10.1099/mgen.0.000211.

26 Zankari Ea, Hasman Henrik, Cosentino Salvatore, Vestergaard Martin, Rasmussen Simon, Lund Ole, et al. Identification of acquired antimicrobial resistance genes. J Antimicrob Chemother 2012;67(11):2640–4. Doi: 10.1093/jac/dks261.

27 Joensen Katrine Grimstrup, Scheutz Flemming, Lund Ole, Hasman Henrik, Kaas Rolf S., Nielsen Eva M., et al. Real-time whole-genome sequencing for routine typing, surveillance, and outbreak detection of verotoxigenic Escherichia coli. J Clin Microbiol 2014;52(5):1501–10. Doi: 10.1128/JCM.03617-13.

28 Chen Lihong, Zheng Dandan, Liu Bo, Yang Jian, Jin Qi. VFDB 2016: hierarchical and refined dataset for big data analysis--10 years on. Nucleic Acids Res 2016;44(D1):D694–697. Doi: 10.1093/nar/gkv1239.

29 Clermont Olivier, Condamine Bénédicte, Dion Sara, Gordon David M., Denamur Erick. The E phylogroup of Escherichia coli is highly diverse and mimics the whole E. coli species population structure. Environ Microbiol 2021;23(11):7139–51. Doi: 10.1111/1462-2920.15742.

30 Page Andrew J., Cummins Carla A., Hunt Martin, Wong Vanessa K., Reuter Sandra, Holden Matthew T. G., et al. Roary: rapid large-scale prokaryote pan genome analysis. Bioinforma Oxf Engl 2015;31(22):3691–3. Doi: 10.1093/bioinformatics/btv421.

31 Minh Bui Quang, Schmidt Heiko A., Chernomor Olga, Schrempf Dominik, Woodhams Michael D., von Haeseler Arndt, et al. IQ-TREE 2: New Models and Efficient Methods for Phylogenetic Inference in the Genomic Era. Mol Biol Evol 2020;37(5):1530–4. Doi: 10.1093/molbev/msaa015.

32 Letunic Ivica, Bork Peer. Interactive Tree Of Life (iTOL) v5: an online tool for phylogenetic tree display and annotation. Nucleic Acids Res 2021;49(W1):W293–6. Doi: 10.1093/nar/gkab301.

33 Berlyn Mary K. B. Linkage Map of Escherichia coli K-12, Edition 10: The Traditional Map. Microbiol Mol Biol Rev 1998;62(3):814–984.

34 Li Heng. Minimap2: pairwise alignment for nucleotide sequences. Bioinforma Oxf Engl 2018;34(18):3094–100. Doi: 10.1093/bioinformatics/bty191.

35 Vaser Robert, Sović Ivan, Nagarajan Niranjan, Šikić Mile. Fast and accurate de novo genome assembly from long uncorrected reads. Genome Res 2017;27(5):737–46. Doi: 10.1101/gr.214270.116.

36 Wick Ryan R., Holt Kathryn E. Benchmarking of long-read assemblers for prokaryote whole genome sequencing. F1000Research 2019;8:2138. Doi: 10.12688/f1000research.21782.4.

37 Wick Ryan R., Judd Louise M., Gorrie Claire L., Holt Kathryn E. Unicycler: Resolving bacterial genome assemblies from short and long sequencing reads. PLoS Comput Biol 2017;13(6):e1005595. Doi: 10.1371/journal.pcbi.1005595.

38 Alikhan Nabil-Fareed, Petty Nicola K., Ben Zakour Nouri L., Beatson Scott A. BLAST Ring Image Generator (BRIG): simple prokaryote genome comparisons. BMC Genomics 2011;12:402. Doi: 10.1186/1471-2164-12-402.

39 Gu Zuguang, Gu Lei, Eils Roland, Schlesner Matthias, Brors Benedikt. circlize Implements and enhances circular visualization in R. Bioinforma Oxf Engl 2014;30(19):2811–2. Doi: 10.1093/bioinformatics/btu393.

40 Brettin Thomas, Davis James J., Disz Terry, Edwards Robert A., Gerdes Svetlana, Olsen Gary J., et al. RASTtk: a modular and extensible implementation of the RAST algorithm for building custom annotation pipelines and annotating batches of genomes. Sci Rep 2015;5:8365. Doi: 10.1038/srep08365.

41 Valens Michèle, Penaud Stéphanie, Rossignol Michèle, Cornet François, Boccard Frédéric. Macrodomain organization of the Escherichia coli chromosome. EMBO J 2004;23(21):4330–41. Doi: 10.1038/sj.emboj.7600434.

42 Diard Médéric, Garry Louis, Selva Marjorie, Mosser Thomas, Denamur Erick, Matic Ivan. Pathogenicity-associated islands in extraintestinal pathogenic Escherichia coli are fitness elements involved in intestinal colonization. J Bacteriol 2010;192(19):4885–93. Doi: 10.1128/JB.00804-10.

43 Hazen Tracy H., Sahl Jason W., Fraser Claire M., Donnenberg Michael S., Scheutz Flemming, Rasko David A. Refining the pathovar paradigm via phylogenomics of the attaching and effacing Escherichia coli. Proc Natl Acad Sci 2013;110(31):12810–5. Doi: 10.1073/pnas.1306836110.

44 Moremi Nyambura, Claus Heike, Vogel Ulrich, Mshana Stephen E. Faecal carriage of CTX-M extended-spectrum beta-lactamase-producing Enterobacteriaceae among street children dwelling in Mwanza city, Tanzania. PloS One 2017;12(9):e0184592. Doi: 10.1371/journal.pone.0184592.

45 Bailey Jannine K., Pinyon Jeremy L., Anantham Sashindran, Hall Ruth M. Distribution of human commensal Escherichia coli phylogenetic groups. J Clin Microbiol 2010;48(9):3455–6. Doi: 10.1128/JCM.00760-10.

46 Roer Louise, Overballe-Petersen Søren, Hansen Frank, Schønning Kristian, Wang Mikala, Røder Bent L., et al. Escherichia coli Sequence Type 410 Is Causing New International High-Risk Clones. MSphere 2018;3(4):e00337–18. Doi: 10.1128/mSphere.00337-18.

47 Armand-Lefèvre Laurence, Rondinaud Emilie, Desvillechabrol Dimitri, Mullaert Jimmy, Clermont Olivier, Petitjean Marie, et al. Dynamics of extended-spectrum beta-lactamase-producing Enterobacterales colonization in long-term carriers following travel abroad. Microb Genomics n.d.;7(7):000576. Doi: 10.1099/mgen.0.000576.

48 Ingle Danielle J., Levine Myron M., Kotloff Karen L., Holt Kathryn E., Robins-Browne Roy M. Dynamics of antimicrobial resistance in intestinal Escherichia coli from children in community settings in South Asia and sub-Saharan Africa. Nat Microbiol 2018;3(9):1063–73. Doi: 10.1038/s41564-018-0217-4.

49 Hernandes Rodrigo T., Hazen Tracy H., dos Santos Luís F., Richter Taylor K. S., Michalski Jane M., Rasko David A. Comparative genomic analysis provides insight into the phylogeny and virulence of atypical enteropathogenic Escherichia coli strains from Brazil. PLoS Negl Trop Dis 2020;14(6):e0008373. Doi: 10.1371/journal.pntd.0008373.

50 Massot Méril, Daubié Anne-Sophie, Clermont Olivier, Jauréguy Françoise, Couffignal Camille, Dahbi Ghizlane, et al. Phylogenetic, virulence and antibiotic resistance characteristics of commensal strain populations of Escherichia coli from community subjects in the Paris area in 2010 and evolution over 30 years. Microbiol Read Engl 2016;162(4):642–50. Doi: 10.1099/mic.0.000242.

51 Nicolas-Chanoine Marie-Hélène, Bertrand Xavier, Madec Jean-Yves. Escherichia coli ST131, an intriguing clonal group. Clin Microbiol Rev 2014;27(3):543–74. Doi: 10.1128/CMR.00125-13.

52 Milenkov Milen, Rasoanandrasana Saida, Rahajamanana Lalaina Vonintsoa, Rakotomalala Rivo Solo, Razafindrakoto Catherine Ainamalala, Rafalimanana Christian, et al. Prevalence, Risk Factors, and Genetic Characterization of Extended-Spectrum Beta-Lactamase Escherichia coli Isolated From Healthy Pregnant Women in Madagascar. Front Microbiol 2021;12:786146. Doi: 10.3389/fmicb.2021.786146.

53 Lavollay M., Mamlouk K., Frank T., Akpabie A., Burghoffer B., Ben Redjeb S., et al. Clonal dissemination of a CTX-M-15 beta-lactamase-producing Escherichia coli strain in the Paris area, Tunis, and Bangui. Antimicrob Agents Chemother 2006;50(7):2433–8. Doi: 10.1128/AAC.00150-06.

54 Rodríguez I., Thomas K., Van Essen A., Schink A.-K., Day M., Chattaway M., et al. Chromosomal location of blaCTX-M genes in clinical isolates of Escherichia coli from Germany, The Netherlands and the UK. Int J Antimicrob Agents 2014;43(6):553–7. Doi: 10.1016/j.ijantimicag.2014.02.019.

55 Shawa Misheck, Furuta Yoshikazu, Mulenga Gillan, Mubanga Maron, Mulenga Evans, Zorigt Tuvshinzaya, et al. Novel chromosomal insertions of ISEcp1-blaCTX-M-15 and diverse antimicrobial resistance genes in Zambian clinical isolates of Enterobacter cloacae and Escherichia coli. Antimicrob Resist Infect Control 2021;10(1):79. Doi: 10.1186/s13756-021-00941-8.

56 Zeng Shihan, Luo Jiajun, Li Xiaoyan, Zhuo Chao, Wu Aiwu, Chen Xiankai, et al. Molecular Epidemiology and Characteristics of CTX-M-55 Extended-Spectrum β-Lactamase-Producing Escherichia coli From Guangzhou, China. Front Microbiol 2021;12:3046. Doi: 10.3389/fmicb.2021.730012.

57 Hamamoto Kouta, Ueda Shuhei, Toyosato Takehiko, Yamamoto Yoshimasa, Hirai Itaru. High Prevalence of Chromosomal blaCTX-M-14 in Escherichia coli Isolates Possessing blaCTX-M-14. Antimicrob Agents Chemother 2016;60(4):2582–4. Doi: 10.1128/AAC.00108-16.

58 Hamamoto Kouta, Hirai Itaru. Characterisation of chromosomally-located blaCTX-M and its surrounding sequence in CTX-M-type extended-spectrum β-lactamase-producing Escherichia coli isolates. J Glob Antimicrob Resist 2019;17:53–7. Doi: 10.1016/j.jgar.2018.11.006.

59 Lehtinen Sonja, Huisman Jana S., Bonhoeffer Sebastian. Evolutionary mechanisms that determine which bacterial genes are carried on plasmids. Evol Lett 2021;5(3):290–301. Doi: 10.1002/evl3.226.

60 Rodríguez-Beltrán Jerónimo, DelaFuente Javier, León-Sampedro Ricardo, MacLean R. Craig, San Millán Álvaro. Beyond horizontal gene transfer: the role of plasmids in bacterial evolution. Nat Rev Microbiol 2021;19(6):347–59. Doi: 10.1038/s41579-020-00497-1.

61 Ludden Catherine, Decano Arun Gonzales, Jamrozy Dorota, Pickard Derek, Morris Dearbhaile, Parkhill Julian, et al. Genomic surveillance of Escherichia coli ST131 identifies local expansion and serial replacement of subclones. Microb Genomics n.d.;6(4):e000352. Doi: 10.1099/mgen.0.000352.

62 Massot Méril, Châtre Pierre, Condamine Bénédicte, Métayer Véronique, Clermont Olivier, Madec Jean-Yves, et al. Interplay between Bacterial Clones and Plasmids in the Spread of Antibiotic Resistance Genes in the Gut: Lessons from a Temporal Study in Veal Calves. Appl Environ Microbiol 2021;87(24):e0135821. Doi: 10.1128/AEM.01358-21.

63 Miltgen Guillaume, Martak Daniel, Valot Benoit, Kamus Laure, Garrigos Thomas, Verchere Guillaume, et al. One Health compartmental analysis of ESBL-producing Escherichia coli on Reunion Island reveals partitioning between humans and livestock. J Antimicrob Chemother 2022;77(5):1254–62. Doi: 10.1093/jac/dkac054.

64 Ruppé Etienne, Lixandru Brandusa, Cojocaru Radu, Büke Cagri, Paramythiotou Elisabeth, Angebault Cécile, et al. Relative fecal abundance of extended-spectrum-β-lactamase-producing Escherichia coli strains and their occurrence in urinary tract infections in women. Antimicrob Agents Chemother 2013;57(9):4512–7. Doi: 10.1128/AAC.00238-13.

65 Andersen Christopher T., Langendorf Céline, Garba Souna, Sayinzonga-Makombe Nathan, Mambula Christopher, Mouniaman Isabelle, et al. Risk of community- and hospital-acquired bacteremia and profile of antibiotic resistance in children hospitalized with severe acute malnutrition in Niger. Int J Infect Dis 2022:S1201-9712(22)00184-9. Doi: 10.1016/j.ijid.2022.03.047.

