## Supplementary Figure 1 for "Faecal carriage of ESBL-producing *Escherichia coli* in a remote region of Niger"

**Supplementary Figure 1.** A. Distribution of core genome SNP distances within and between haplogroups. For each isolate, distance in core genome SNPs has been determined. The dotted line represents the cut-off of 120 SNPs used to determine if isolates were considered as belonging to the same clone or not. Vertical bars represent interquartile ranges. B. Distribution of core genome patristic distances within and between haplogroups.

A.

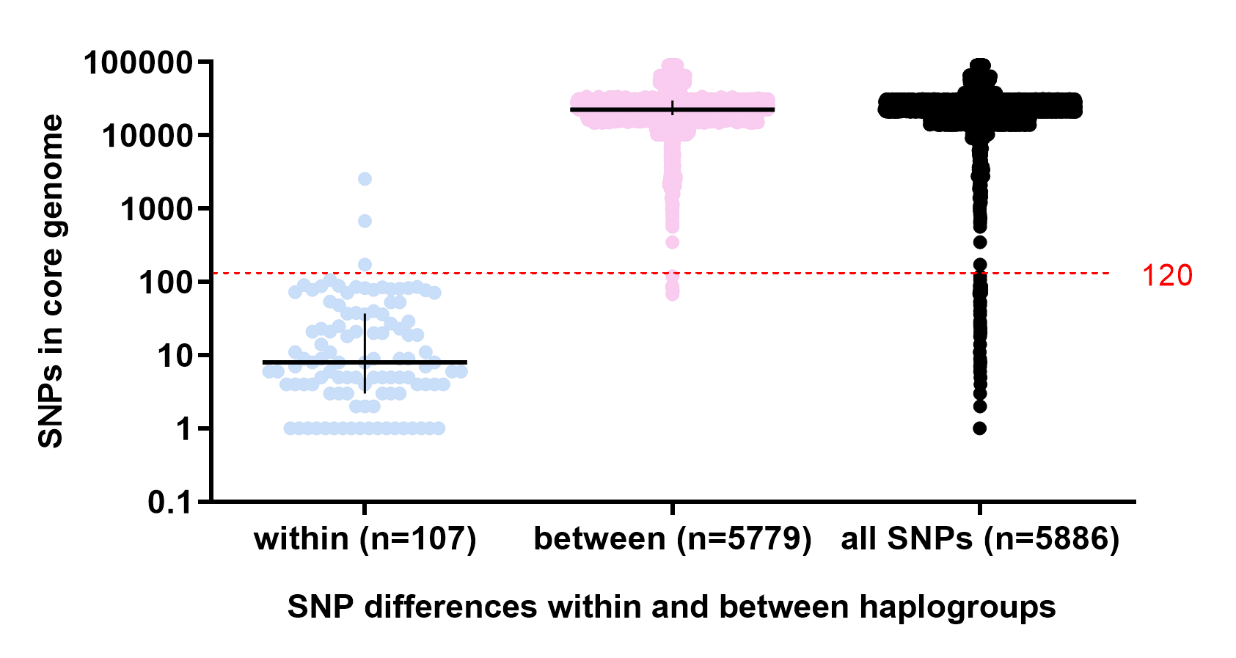

B.

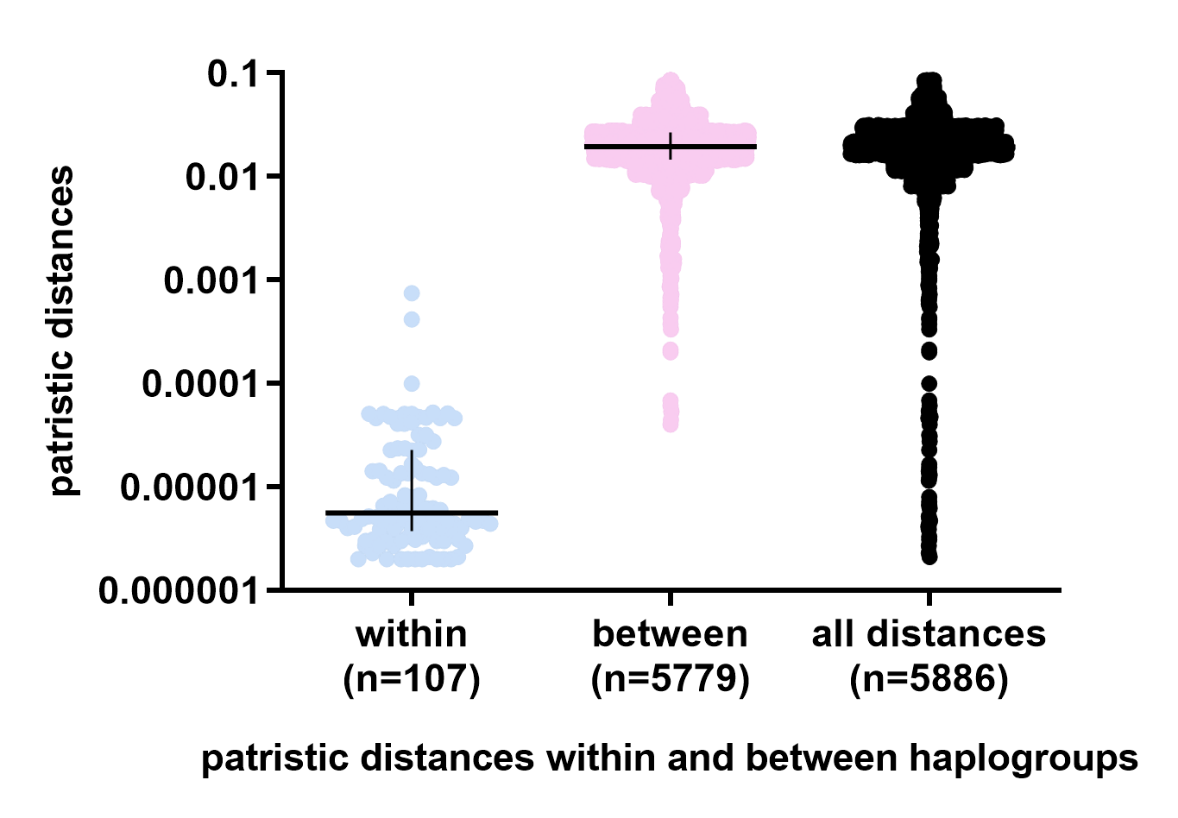
