## Supplementary Figure 2 for "Faecal carriage of ESBL-producing *Escherichia coli* in a remote region of Niger"

**Supplementary Figure 2**. Alignments of plasmids displaying high similarity (less than 1% difference in the nucleotide sequences or ≤2 large insertions or deletions)
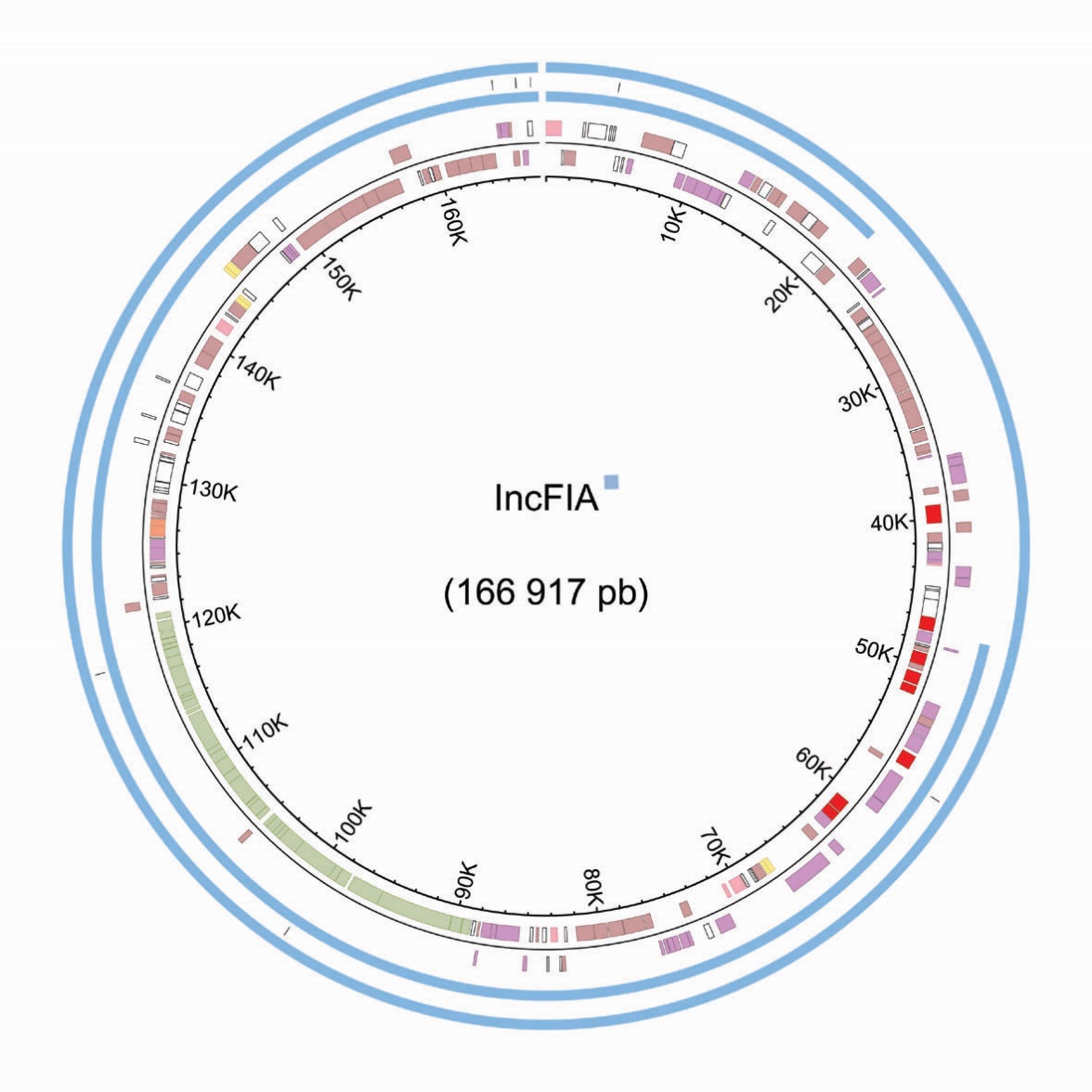

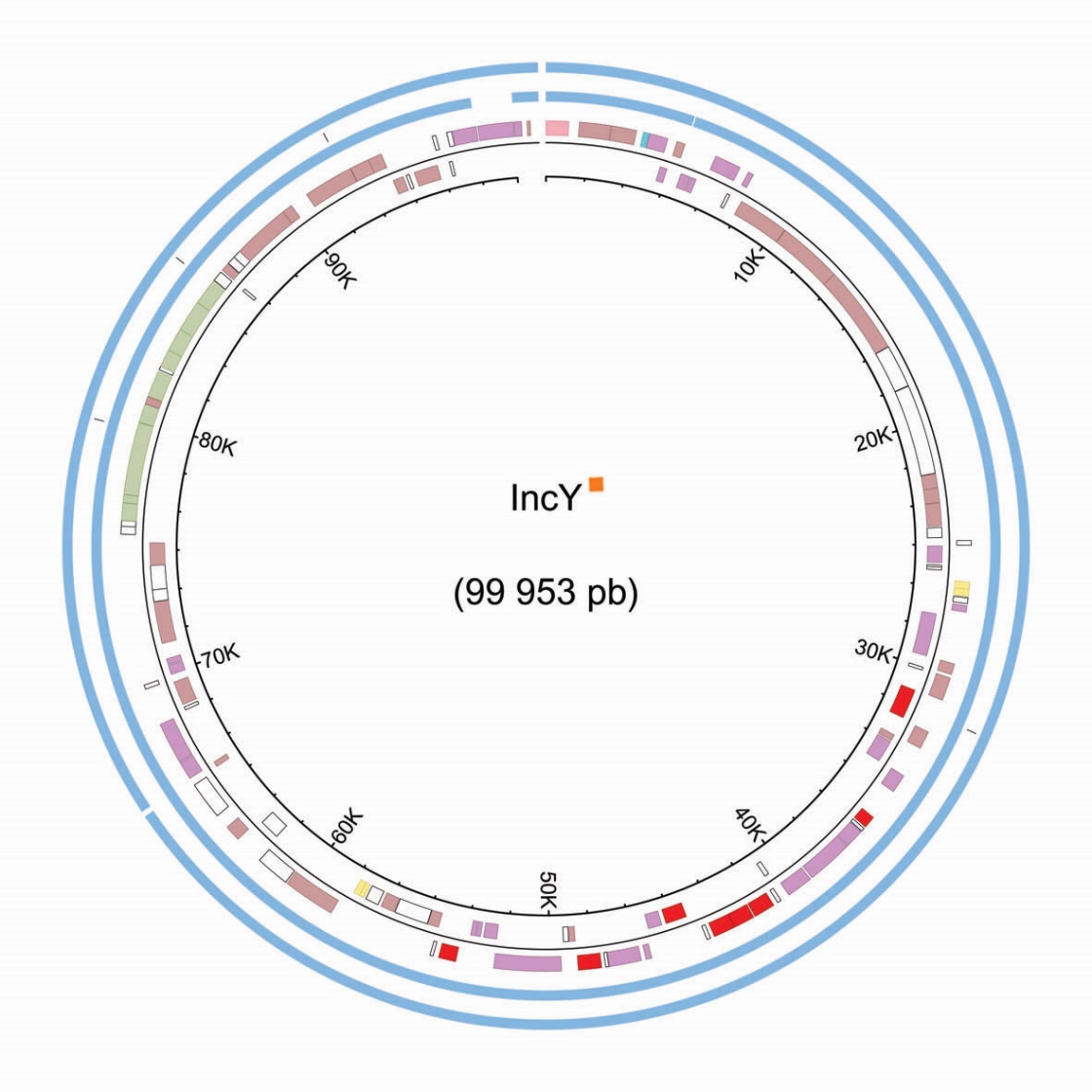

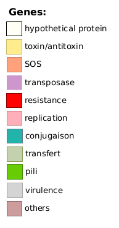

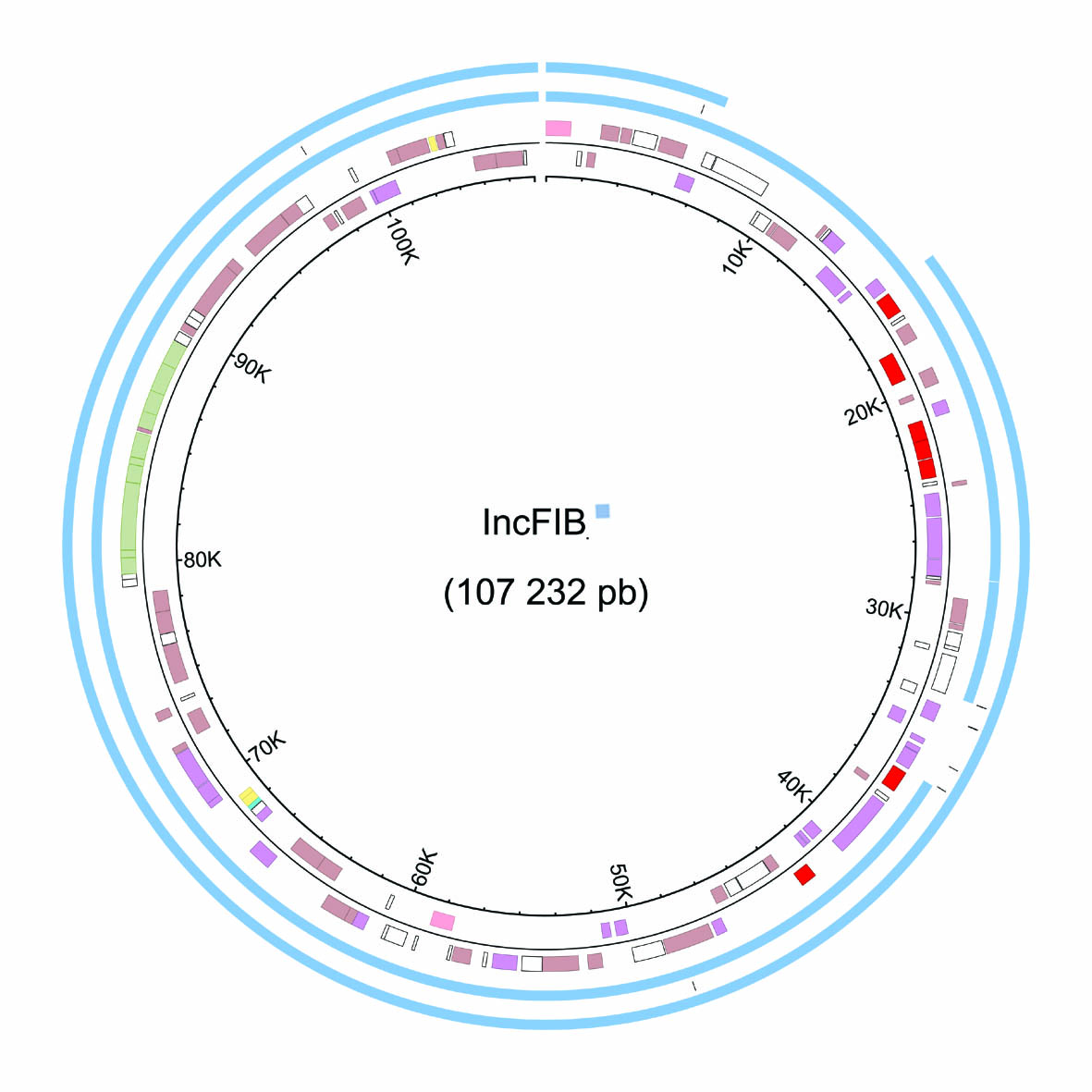

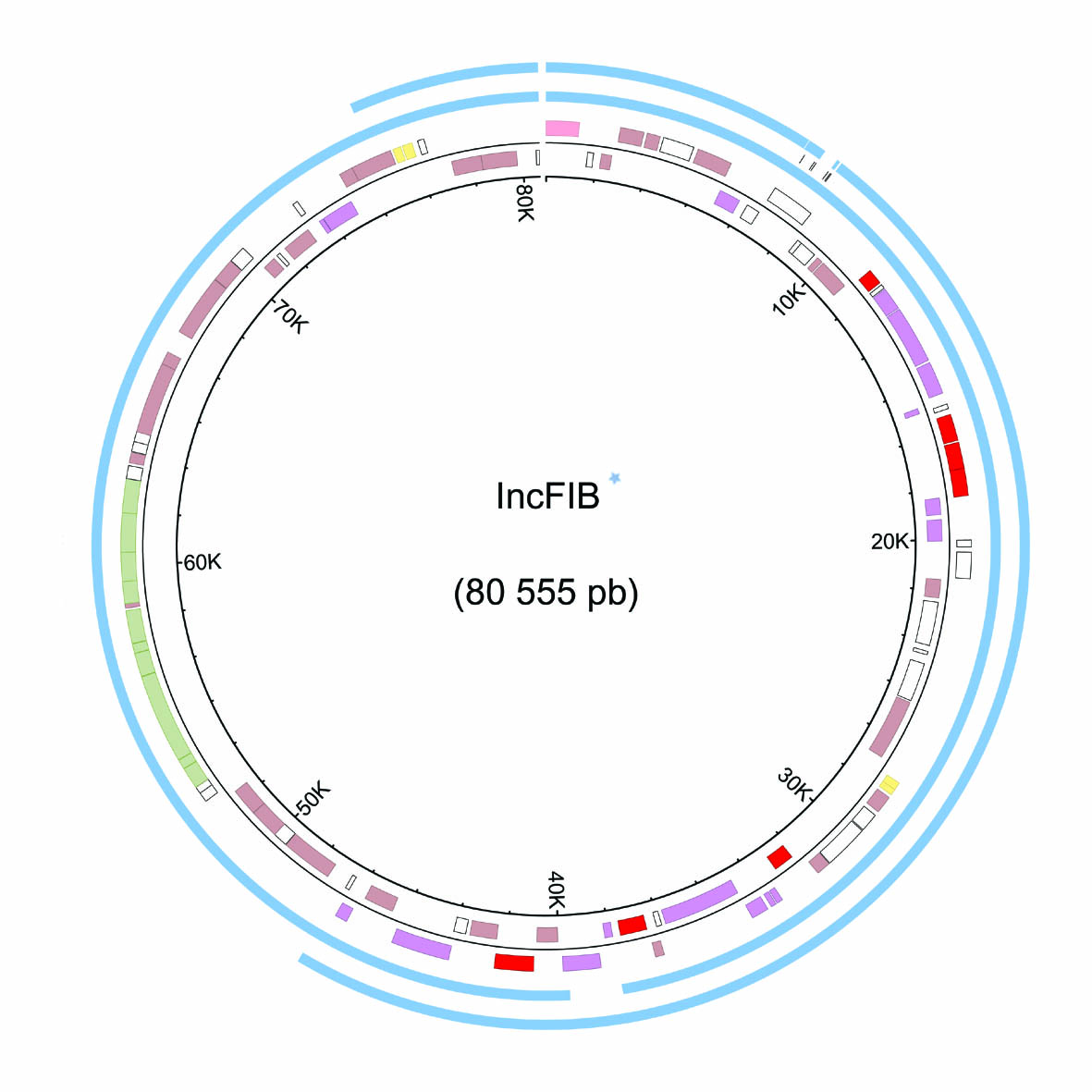

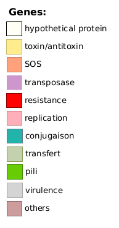

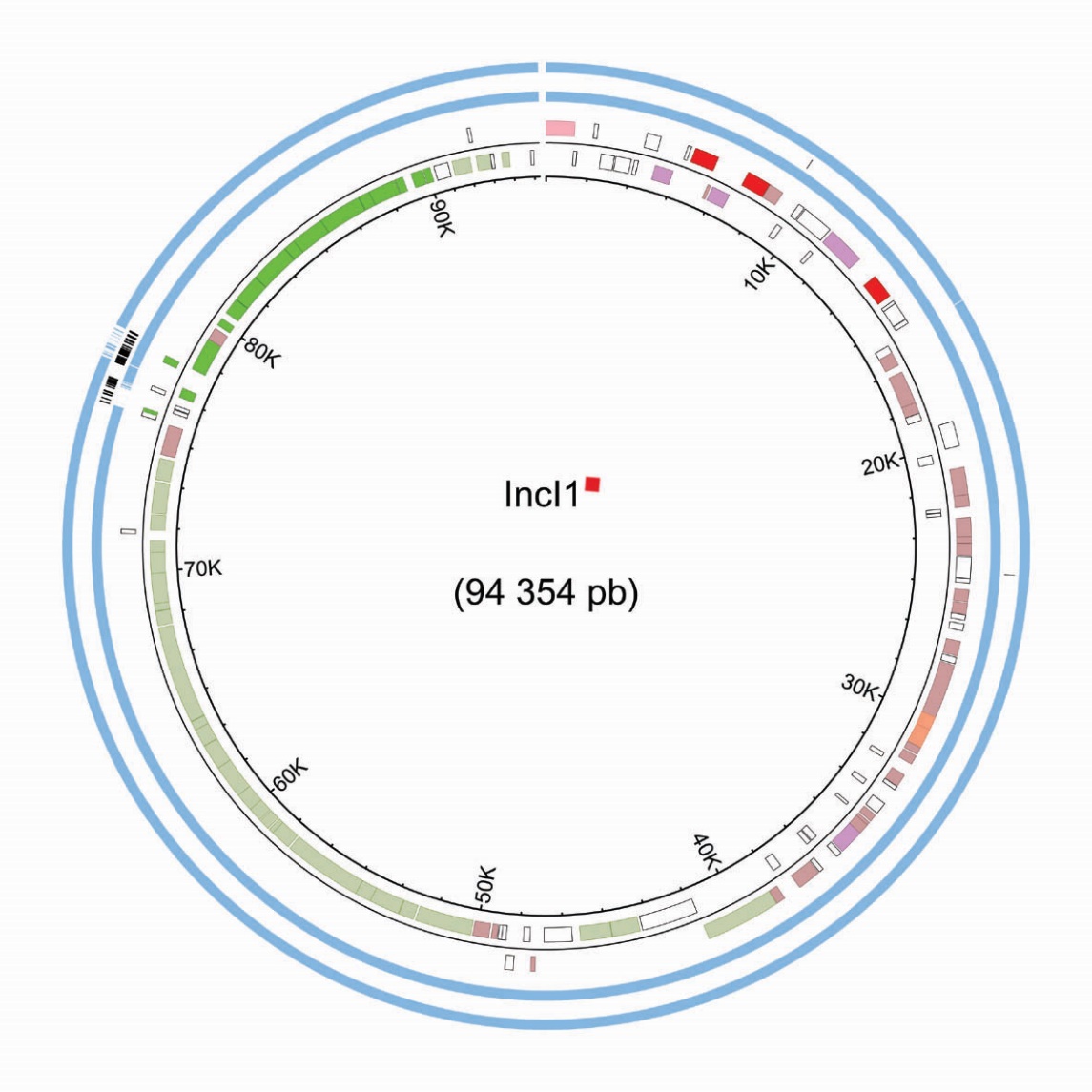

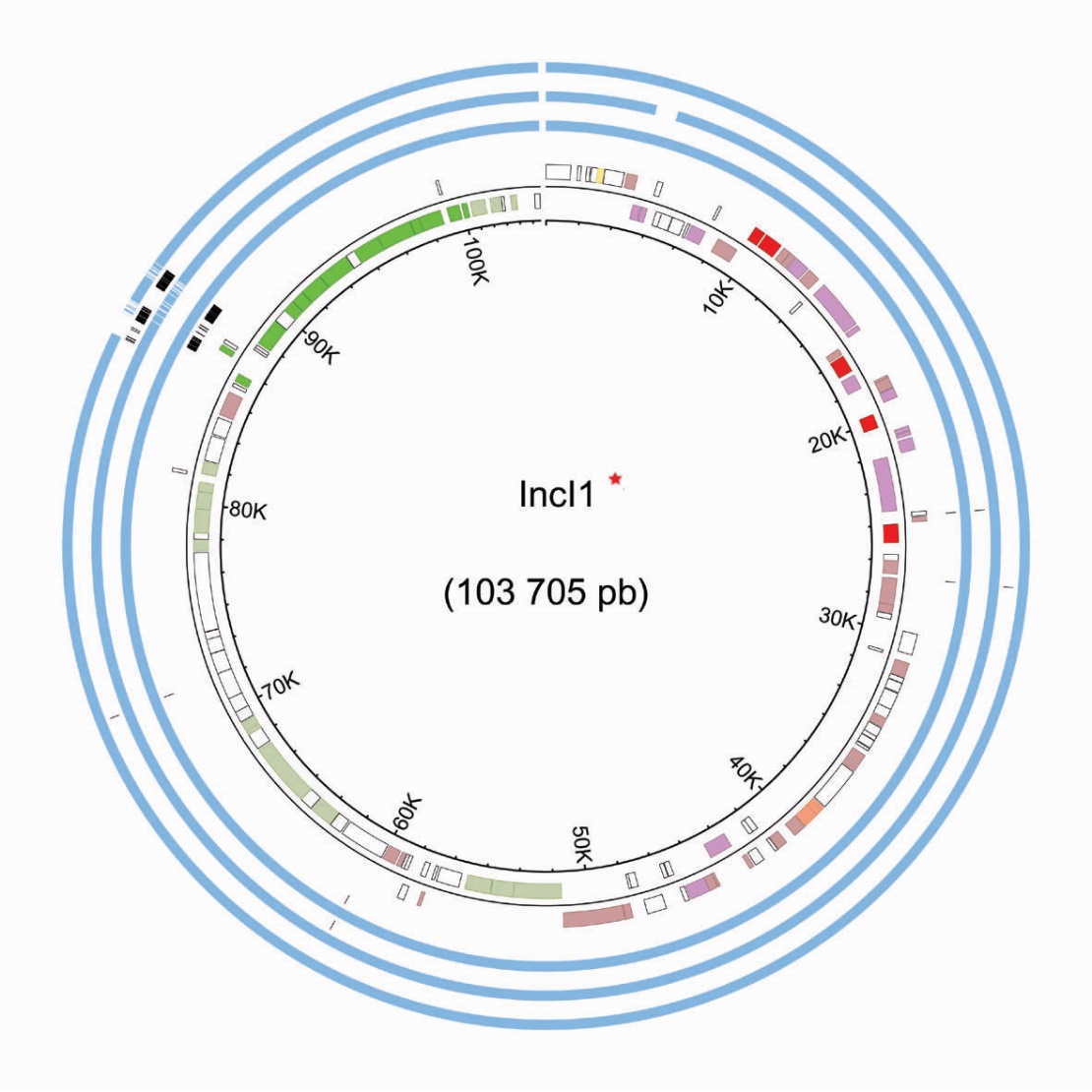

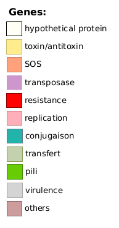
