## Supplementary Table3 for "Faecal carriage of ESBL-producing *Escherichia coli* in a remote region of Niger"

| **Integration site code** | **CDS upstream genetic mobile element** | **CDS downstream genetic mobile element** | **Position on the chromosome of *E. coli* K12 (in minutes)** |
| --- | --- | --- | --- |
| A | Uncharacterized protein YaiT, interrupted by IS3E | Porphobilinogen synthase (hemB) | 8.4 |
| B | ND | Hemolysin E (hlyE) | 26.5 |
| C | serine tRNA | Clp ATP-dependent protease, ATP-binding | 22.2 |
| D | tRNA-Met-CAT | Outer membrane beta-barrel assembly protein BamE | 63.5 |
| E | Diguanylate cyclase ycdT | Diguanylate cyclase ycdT | 23.5 |
| F | Outer membrane protein W precursor YciD | Outer membrane protein W precursor YciD | 28.3 |
| G | Methyl-accepting chemotaxis protein III (ribose and galactose chemoreceptor protein) Trg | Uncharacterized protein YdcA | 32.11 |
| H | Glucuronide transporter UidB | Glucuronide transporter UidB | 36.4 |
| I | HipB protein / Antitoxin HigA | Toxin HigB / Protein kinase domain of HipA | 34.3 |
| J | c-di-GMP phosphodiesterase | c-di-GMP phosphodiesterase | 56.6 |
| K | Inner membrane protein YbhI | Inner membrane protein YbhI | 17.3 |
| L | ABC-type sugar transport system, periplasmic binding protein YcjN | ABC-type sugar transport system, periplasmic binding protein YcjN | 29.5 |
| M | Pyridoxine 4-dehydrogenase PhxI (EC 1.1.1.65)(YdbD) | DUF2773 domain-containing bactofilin YdbC | 31.7 |
| N | Biofilm PGA synthesis auxiliary protein PgaC | Biofilm PGA synthesis auxiliary protein PgaD | 23.4 |
| O | Uncharacterized GST-like protein yncG | L-asparagine permease ansP | 32.9 |
| P | Glutamate racemase | Ethanolamine utilization protein EutA | 55.3 |
| Q | ND | Tyramine oxidase TynA | 31.2 |
| R | Putative inner membrane protein YqgA | ND | 67.0 |
| S | Guanine-hypoxanthine permease | ND | 65.2 |

**Supplementary Table 3**. Correspondence between integration site code and CDS of *E. coli* K12 chromosome located upstream and downstream *bla*_CTX-M-15_ region_._
